# ProtGPT3: an Open-source family of Promptable and Aligned Protein Language Models

**DOI:** 10.64898/2026.06.04.730041

**Authors:** Michele Garibbo, Gerard Boxó, Filippo Stocco, Ramiro Illanes-Vicioso, Lasse Middendorf, Noelia Ferruz

**Affiliations:** Centre for Genomic Regulation (CRG), Barcelona Institute of Science and Technology, Barcelona, Spain; Universitat Pompeu Fabra (UPF), Barcelona, Spain

## Abstract

Generative protein language models (pLMs) enable exploration of vast sequence spaces for protein design, but reliably controlling generation toward desired functional families remains challenging. While protein generation has broadly followed trends in NLP, two directions remain underexplored: alignment methods that optimize model behavior toward design objectives, and prompting-based control at inference time without fine-tuning. We introduce ProtGPT3, an open-source family of protein language models spanning 112M to 10B parameters and integrated with the Hugging Face ecosystem. The suite includes both single-sequence and multiple sequence alignment (MSA)-promptable models, enabling flexible conditioning for generation. Across model scales and protein families, we systematically compare supervised fine-tuning and few-shot prompting using homologous sequences. Analogous to how large language models (LLMs) are routinely aligned with user intent, we study post-training alignment in single-sequence models using sequence-complexity and structure-confidence metrics across the proteome. We find that alignment reduces low-complexity generations while preserving sequence diversity. Furthermore, we show that few-shot prompting is a competitive and more scalable alternative to supervised fine-tuning for controlled generation. In a low-data defluorinase case study, ProtGPT3-MSA achieved higher computational success rates than fine-tuned baselines and produced designs that were soluble and expressed following experimental validation. Finally, we explore the potential of inference-time compute in MSA models by introducing a homolog-based Feynman–Kac inference procedure for steering protein generation toward desired targets. All models are publicly available at https://huggingface.co/collections/AI4PD/protgpt3-family.

## 1 Introduction

Proteins found in nature represent a vanishingly small sample of the astronomical number of possible sequences that can be formed. Natural proteins perform a remarkable diversity of functions, from catalyzing reactions to regulating cellular pathways and enabling immune responses. Yet their properties are shaped by evolutionary pressures that rarely align with industrial or therapeutic requirements. The design of artificial proteins with novel and customizable functions has therefore been a central goal of biochemistry. Historically, this was viewed primarily as a basic research endeavor: a stringent test of our understanding of protein folding and molecular interactions. Over the last five years, however, protein design has evolved into an applied discipline of broad technological relevance, driven by advances in deep learning, structure prediction, and the rapid expansion of biological sequence databases [24].

Protein language models (pLMs) have emerged as versatile tools for protein science, supporting tasks ranging from structure and fitness prediction [27] to *de novo* sequence generation [55]. In particular, autoregressive pLMs have shown strong generative capabilities, enabling the design of diverse proteins, including experimentally validated enzymes and fluorescent proteins, often with sequence identities as low as 30% to known natural proteins [31]. While the development of pLMs has mirrored advances in NLP, several approaches routinely used for large language models (LLMs) remain underexplored in protein design. First, modern LLMs are routinely post-trained to *align* them with user intent, with the goal of making them more helpful, truthful, and harmless, using reinforcement learning (RL) and preference-optimization methods [38]. Although RL is emerging as a strategy for aligning generative pLMs for protein design [50], studies of pLM alignment at the whole-proteome level remain limited. Second, following GPT-2, LLMs increasingly shifted from task-specific fine-tuning toward in-context or few-shot prompting [6, 37], in which a model conditions on a small number of examples at inference time without updating its weights. In protein engineering, by contrast, users often rely on fine-tuning comparatively small pLMs on curated task-specific datasets. While fine-tuning approaches have achieved considerable success [29, 43, 42], they present two key limitations: (1) it can be difficult to determine the appropriate extent of fine-tuning for a new downstream task without overwriting or degrading previously acquired knowledge, thereby reducing the model’s ability to generalize across tasks; and (2) they shift the computational burden of adaptation onto end users.

Here, we introduce ProtGPT3, a family of open-source autoregressive pLMs spanning 112M to 10B parameters and trained with size-optimized data mixtures. We further study post-training alignment of single-sequence models toward desirable protein-generation properties, including high predicted structural confidence and reduced low-complexity content. In parallel, we introduce ProtGPT3-MSA, a homolog-conditioned model that processes multiple related sequences simultaneously and enables inference-time adaptation through prompting rather than task-specific fine-tuning. This shifts the typical workflow in protein design: instead of fine-tuning a separate model for each new family or campaign, users can steer generation from a small set of homologous sequences while preserving the behavior of the underlying pretrained model. Across held-out protein families, prompting ProtGPT3-MSA with as few as 15 homologs produces family-consistent generations and compares favorably to supervised fine-tuning of single-sequence models, particularly in low-data regimes. We further show that prompt composition and Feynman–Kac-style sequential Monte Carlo inference can bias generation toward desired sequence-level objectives without updating model parameters. In an extreme low-data defluorinase case study using only seven experimentally annotated sequences, ProtGPT3-MSA achieves substantially higher computational pass rates than fine-tuned single-sequence baselines and generates designs that are successfully cloned, expressed, and purified in *E. coli*.

Our main contributions are as follows: 1) We release ProtGPT3, a portable and open-source family of autoregressive protein language models spanning 112M to 10B parameters (at https://huggingface.co/collections/AI4PD/protgpt3-family), together with model checkpoints, configuration files, and code for efficient training. 2) We introduce ProtGPT3-MSA, a homolog-conditioned model that enables few-shot family-conditioned generation without task-specific fine-tuning. 3) We provide a systematic comparison between prompt-based conditional generation and supervised fine-tuning with wet-lab validation and 4) we study two complementary strategies for controllable generation: post-training alignment, and inference-time steering.

## 2 Related Work

### Scaling and open sourcing

Scaling model size and training data has been central to improving both LLMs [6, 22, 17] and pLMs. Early generative pLM scaling studies such as RITA released autoregressive models up to 1.2B parameters trained on UniRef100 [16]. Subsequent work explored scaling laws and compute-optimal training for masked and causal pLMs, releasing models up to 7B parameters [10, 45]. The ProGen family extended this direction for protein design: ProGen2 scaled autoregressive pLMs to 6.4B parameters, while ProGen3 introduced sparse models up to 46B parameters and studied how scale affects diversity, expression, and alignment [32, 4]. More recently, xTrimoPGLM explored unified masked and autoregressive pretraining at 100B scale [9], and the Dayhoff Atlas released large-scale protein datasets and models from 117M to 3B parameters [60]. Beyond scaling, usability and openness remain essential for reproducible protein design, motivating our public HuggingFace release with training and inference support.

### Alignment

Scaling pLMs improves representations and generation quality, but does not inherently align models with design objectives such as activity, binding affinity, or stability [17, 10, 8, 4, 50]. We use RL-alignment to describe parameter-updating methods that steer pretrained generative priors toward task objectives using feedback, in contrast to parameter-free inference-time steering [50]. Prior work has therefore focused on aligning pLMs with specific design objectives: ESM-IF1 is aligned with stability using DPO [59], ProGen3 is aligned with activity, binding, and stability signals from ProteinGym and Megascale via IRPO [4], and PPO- or DPO-based approaches have been used for fluorescence optimization and EGFR binder engineering using experimental feedback [5, 49]. In contrast, we focus on a more general and widely observed bias in unconditional generation, low-complexity sequence collapse [4], and study alignment as a mechanism to mitigate this proteome-scale sampling bias rather than optimizing for specific biochemical properties.

### Few-shot family-conditioned generations

Recent protein generative models condition on homologous sequences as in-context examples of a target family, analogous to prompting in LLMs. Early approaches relied on aligned MSAs: MSA Transformer introduced row–column attention over homolog alignments [40], later enabling enzyme design applications [21]. More recent methods use unaligned homolog sets: PoET represents families as order-invariant sequence sets for retrieval-augmented generation [51], while ProtMamba and Bio-xLSTM use long-context sequence models for homology-aware generation [46, 44]. ProFam evaluates whether generated families preserve natural conservation and covariance statistics suitable for AlphaFold2 inputs [58], and Dayhoff unifies single-sequence, aligned-, and unaligned-homolog conditioning, reporting experimental expression rates of 35% [60]. While these works demonstrate the potential of prompting protein language models with homologous sequences for target family generation, to our knowledge, a systematic comparison between this approach and the more traditional paradigm of supervised fine-tuning remains lacking. In ProtGPT3, we address this gap by providing a systematic comparison between prompting and supervised fine-tuning across several target protein families and model scales, including experimental validation for a low-data defluorinase case study.

## 3 Large-Scale Pretraining of a Family of Single-Sequence pLMs

### 3.1 Model architectures

ProtGPT3 is a family of autoregressive decoder-only protein language models trained with a Mixtral-style (sparse) mixture-of-experts (MoE) architecture [20], with 8 experts and the load-balancing scheme. This *sparse* architecture was chosen after comparing its training and inference performance against a popular *dense* large language model architecture (i.e., Llama3[15]) for autoregressive protein prediction. We found the Mixtral architecture achieves marginally worse validation perplexity, while enabling faster training (≈1*/*3 faster) and inference (≈1*/*2 faster), providing the optimal compute approach (see Appendix A.1). These findings are consistent with previous work in autoregressive pLMs [4] and more broadly in the NLP literature [e.g., 28, 20].

Based on Mixtral architecture, we trained three single-sequence models: ProtGPT3-112M, ProtGPT3-1.3B, and ProtGPT3-10B, as well as a multiple sequence alignment (MSA) model, ProtGPT3-MSA (112M). All our models were trained to perform causal language modeling in both directions [e.g., 32, 29, 4], enabling protein generation from either the N-to-C or C-to-N terminal with the help of two special directional tokens encoding the direction at the start of each protein (Appendix (A.2)). ProtGPT3-MSA was trained auto-regressively to predict inputs of 16 concatenated protein sequences one token at a time (up to a max of 16384 tokens), starting from ProtGPT3-112M last training checkpoint. For each input, the 16 protein sequences were sampled from the same MSA (see Dataset 3.2) and separated by a special separator token. We enforced the 16 protein sequence constraint on all training MSAs, even when the context length allowed to fit more sequences to prevent having unbalanced numbers of sequences across training MSAs. Following a similar approach to Yang et al. [60], ProtGPT3-MSA was trained with two main modalities (1) *aligned*: the 16 input sequences were passed to the model aligned including gap tokens (i.e., “ − “) and (2) *unaligned*: the 16 input sequences were passed in full (i.e., including insertions and without gaps). We introduced two extra special tokens at the start of each concatenated set of sequences to specify the modality to the model: *< gap >* denoted the *aligned* modality, while *< no*_*gap >* denoted the *unaligned* one. For maximal training efficiency, all our models were trained with online mini-batch packing with Flash-Attention [25].

### 3.2 Datasets

To train our model family, we leveraged the UniRef90 [2] dataset and the Dayhoff Atlas [60], which at the time of training provided the largest curated public protein dataset (3.34 billion sequences). From the Atlas, we used the GigaRef subset, restricting to non-singleton clusters to ensure higher-quality sequences. This subset comprises 1.89 billion sequences across 243 million clusters at 50% sequence identity. In order to sample training sequences across clusters of varying sizes we adopted a two step sampling procedure, which was shown to achieve the best validation performance on out-of-distribution sequences [4]. First, we sampled a cluster based on its size, following an inverse log law (i.e., 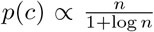 with *n* being the number of sequences in the cluster). Next, we uniformly sample a sequence within the sampled cluster. Prior work suggests that (genomic) UniRef90 sequences are critical for strong pLM performance [10, 60, 30]. Although GigaRef includes some UniRef100-derived sequences, they represent only a small fraction of the data (13M out of 243M clusters), making them relatively underrepresented compared to metagenomic sequences. To prevent metagenomic data from dominating training, we explored two different strategies to preserve the contribution of UniRef sequences. In first strategy, we constructed batches with an equal mix of tokens from GigaRef and UniRef90. In the second strategy, we trained solely on GigaRef but applied a weighted loss to upweight UniRef-derived sequences within it. We evaluated these approaches by training four 112M-parameter models: (1) UniRef90 only, (2) GigaRef only, (3) 50/50 mixed batches, and (4) GigaRef with the weighted loss. For evaluation, we built two validation sets, one based on UniRef90 and another based on GigaRef. We employed MMseqs2 [47] to enforce a maximum of 50% sequence identity between training and validation sequences, while removing any cross contamination by mixing UniRef90 and GigaRef sequences. We found the model trained on 50/50 mixed batches achieved the best performance trade-off across UniRef90 and GigaRef validation perplexity (Fig. 5 in Appendix), resulting in the adapted strategy to train all other models (note the 10B parameter model was trained with 30/70 UniRef90-GigaRef sequences due to an sufficient number of UniRef90 sequences to achieve the optimal number of tokens). All our single-sequence models were trained based on optimal scaling laws for sparse protein language models [4, 10] (Appendix A.2).

ProtGPT3-MSA was trained on 8.5M MSAs from the OpenProteinSet Uniclust30 dataset [1]. From each MSA, 16 sequences were sampled without replacement and concatenated together in random order. A special separator token *< s >* was introduced to mark the sequence boundaries. Each sampled set of 16 sequences was randomly assigned to one of 4 modalities, defined by combinations of generation direction and alignment format. This process was repeated 15 times for each MSA.

### 3.3 Unconditional protein generation across scale

To evaluate the sequence-only models’ performance across scale, we first computed each model’s perplexity on held-out validation sequences that had at most 50% sequence identity to the training set of each model. Importantly, we did this at the protein-cluster level rather than as a single validation average, capturing how uniformly each model covers the diversity of the held-out sequence space. We found a clear effect of scale, with larger models achieving lower perplexity not only on average but across the full distribution of clusters, progressively reducing the tail of sequences for which the smaller models assign high perplexity (Fig 1c & Appendix A.3). In line with previous work [e.g., 4, 10], we compared the models’ generation performance for 1) sequence quality and 2) sequence diversity, by generating 100 sequences using top-p sampling [18] for each possible combination of top-p {0.6, 0.8, 0.9, 1.0} and temperature {0.6, 0.8, 0.9, 1.0} across the 3 model sizes. Sequence quality was measured in terms of predicted pLDDT by ESMFold [27], proportion of terminating sequences (i.e., sequences for which the model correctly produced a terminating EOS token) and proportion of low complexity proteins (i.e., sequences consisting of more than 25 % low complexity regions (LCR)) [14]. Sequence diversity was measured by clustering each model generated sequences at 50% sequence identity and counting the proportion of independent clusters relative to the number of generated sequences [e.g., 10]. We found a clear effect for scale with the largest model (i.e., ProtGPT3-10B) generating the highest quality and most diverse sequences (Fig 1c-d). Importantly, these results are aggregated across all combinations of top-p 0.6, 0.8, 0.9, 1.0 and temperature 0.6, 0.8, 0.9, 1.0, which masks the strong dependence on hyperparameter selection; in practice, specific configurations can yield substantially higher quality than the reported averages. Next, we benchmark all three single-sequence models against ProteinGym [33], again finding that the largest model achieves the best performance (see Table 2 in the Appendix). While our models perform comparably to other single-sequence autoregressive pLMs on substitution DMS datasets, ProtGPT3-10B is among the top 4 performing models on indel DMS datasets (see Table 2 in Appendix).

**Figure 1:**
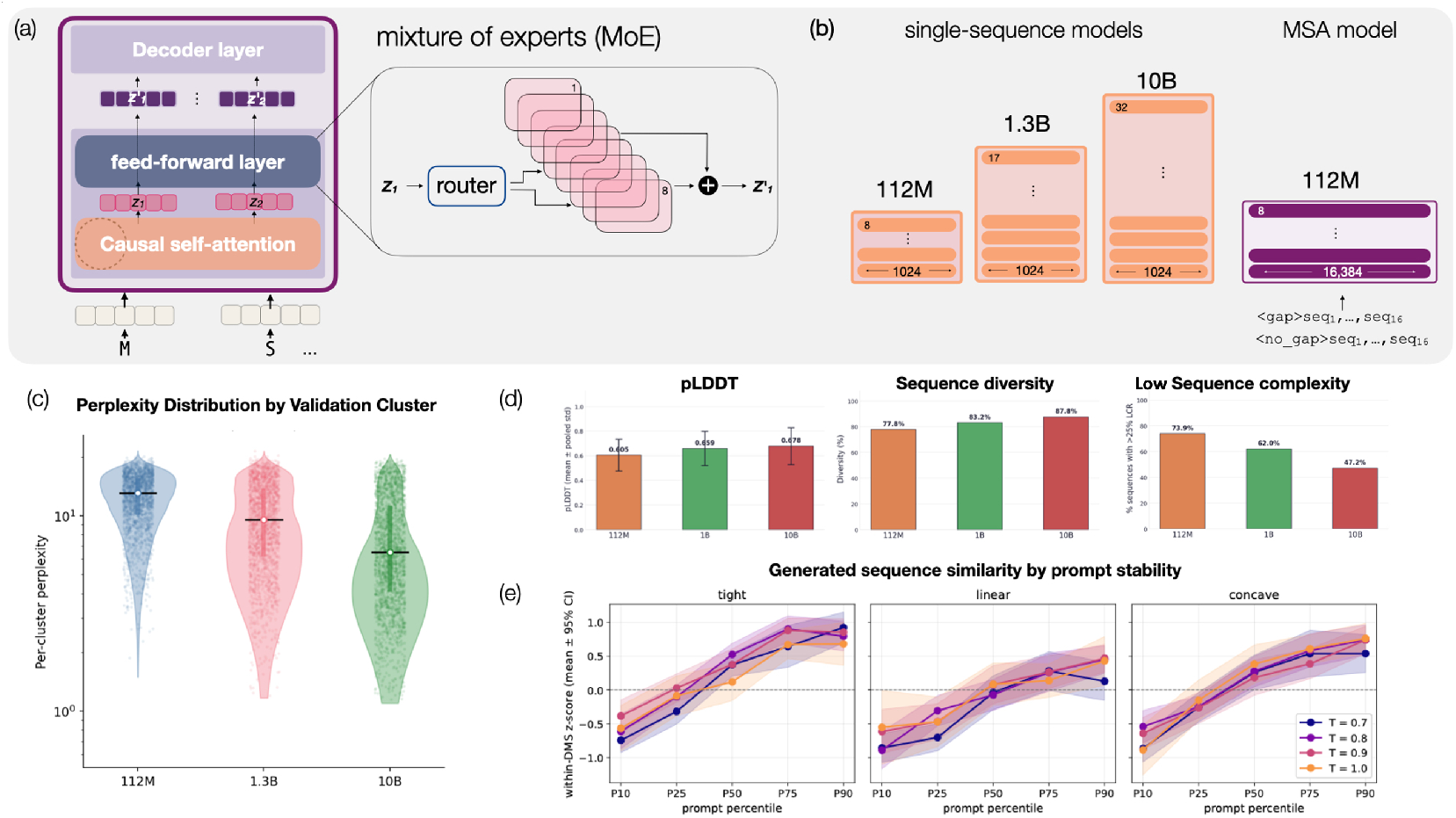
Architecture and pre-training performance of the ProtGPT3 family. **(a)** ProtGPT3 models use a causal Mixtral Mixture-of-Experts architecture. Sequences can be generated in N-to-C or C-to-N direction **(b)** We scaled training of single-sequence models from 112M to 10B parameters. Additionally, we trained ProtGPT3-MSA that can be conditioned on up to 15 aligned or unaligned protein sequences, enabling few-shot prompting for family-conditioning and in-context learning. **(c)** Larger models achieve lower perplexity across the full distribution of held-out validation protein clusters. **(d)** Performance of single-sequence models improves with parameter scale across different metrics. **(e)** In-context learning of protein stability with ProtGPT3-MSA. Prompting ProtGPT3-MSA with more stable protein sequences steers generation towards sequences with higher stability, across different prompting strategies and temperature.

**Figure 2:**
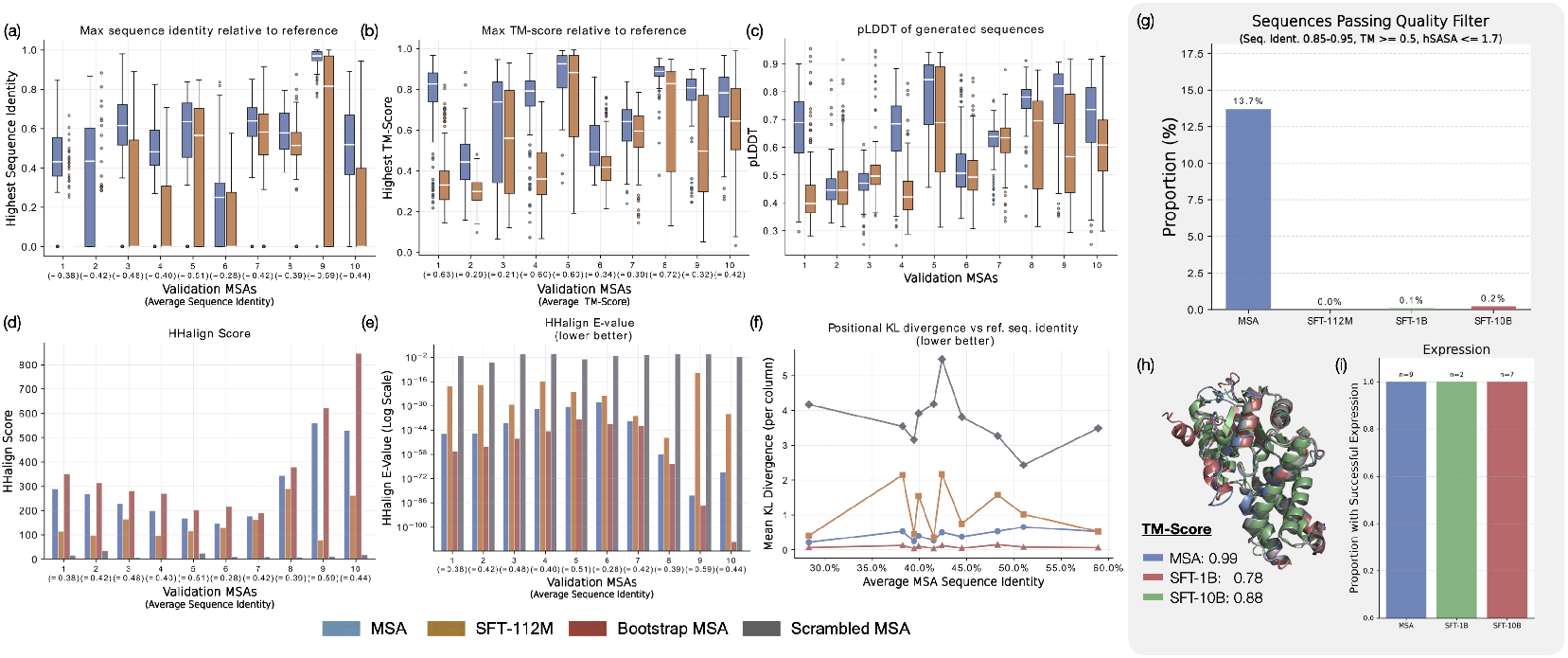
Comparison of MSA Prompting vs. Supervised Fine-Tuning and Defluorinase Design. **(a-f)** Conditional generation comparison prompting ProtGPT3-MSA vs SFT (single-sequence) ProtGPT3-112M for 10 held-out validation MSAs from ProtGPT3-MSA pre-training validation set. See Fig 15 in the Appendix for a comparison with ProtGPT3-10B. **(g)** Computational success rates of designed defluorinases generated with ProtGPT3-MSA and SFT models. **(h)** Superimposed predicted structures (Blue, Green, Red) with highest TM-Score for the respective model towards the experimental solved structure (PDB: 3UMG; Gray). **(i)** Experimental expression rates of successfully cloned designs

**Figure 3:**
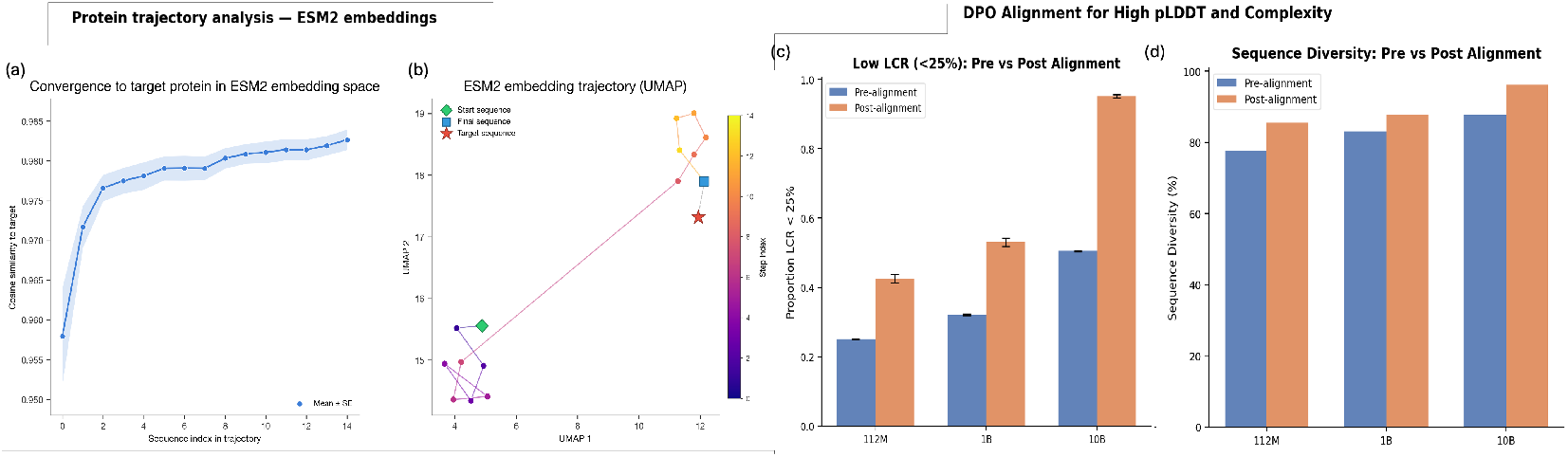
Target generation and inference-time compute and alignment. **(a-b)** At inference time, generation of ProtGPT3-MSA can be steered via Feynman-Kac-style sequential Monte Carlo sampling as demonstrated by traversing ESM2 embedding space between homologous sequences. **(c)** DPO alignment improves sequence complexity across model scale, while **(d)** preserving sequence diversity.

**Figure 4:**
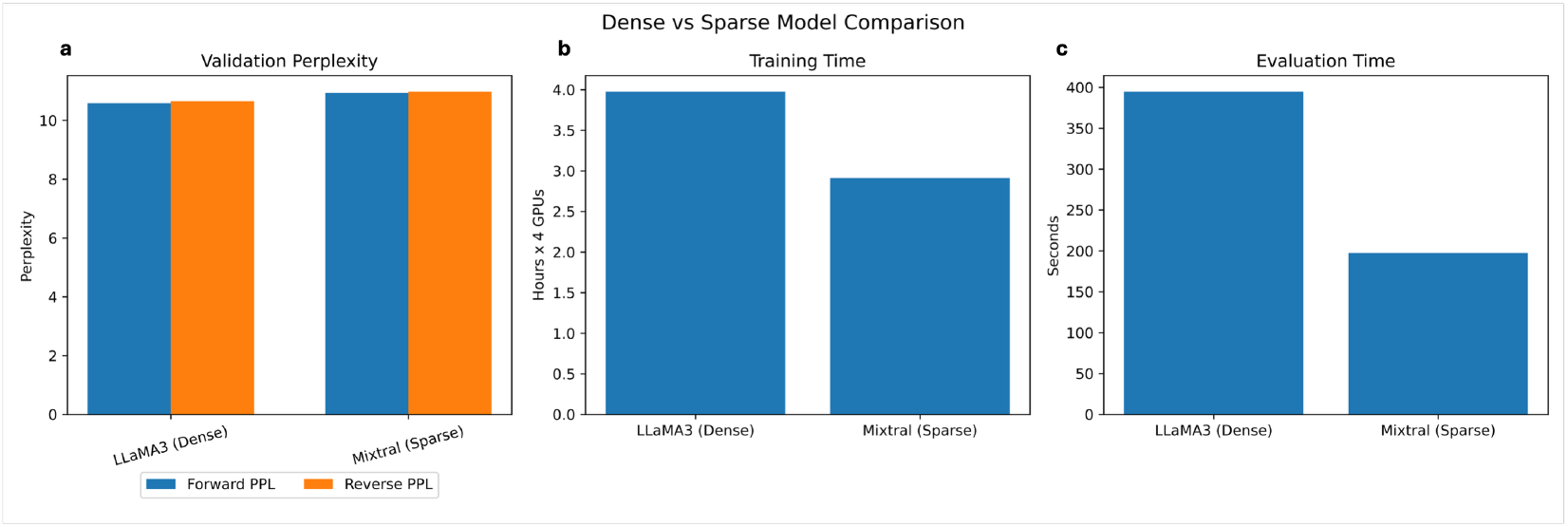
Model architecture: LLama3 (dense) vs Mixtral (sparse) model comparison in terms of a) validation perplexity on held-out protein sequences b) training time and c) inference

**Figure 5:**
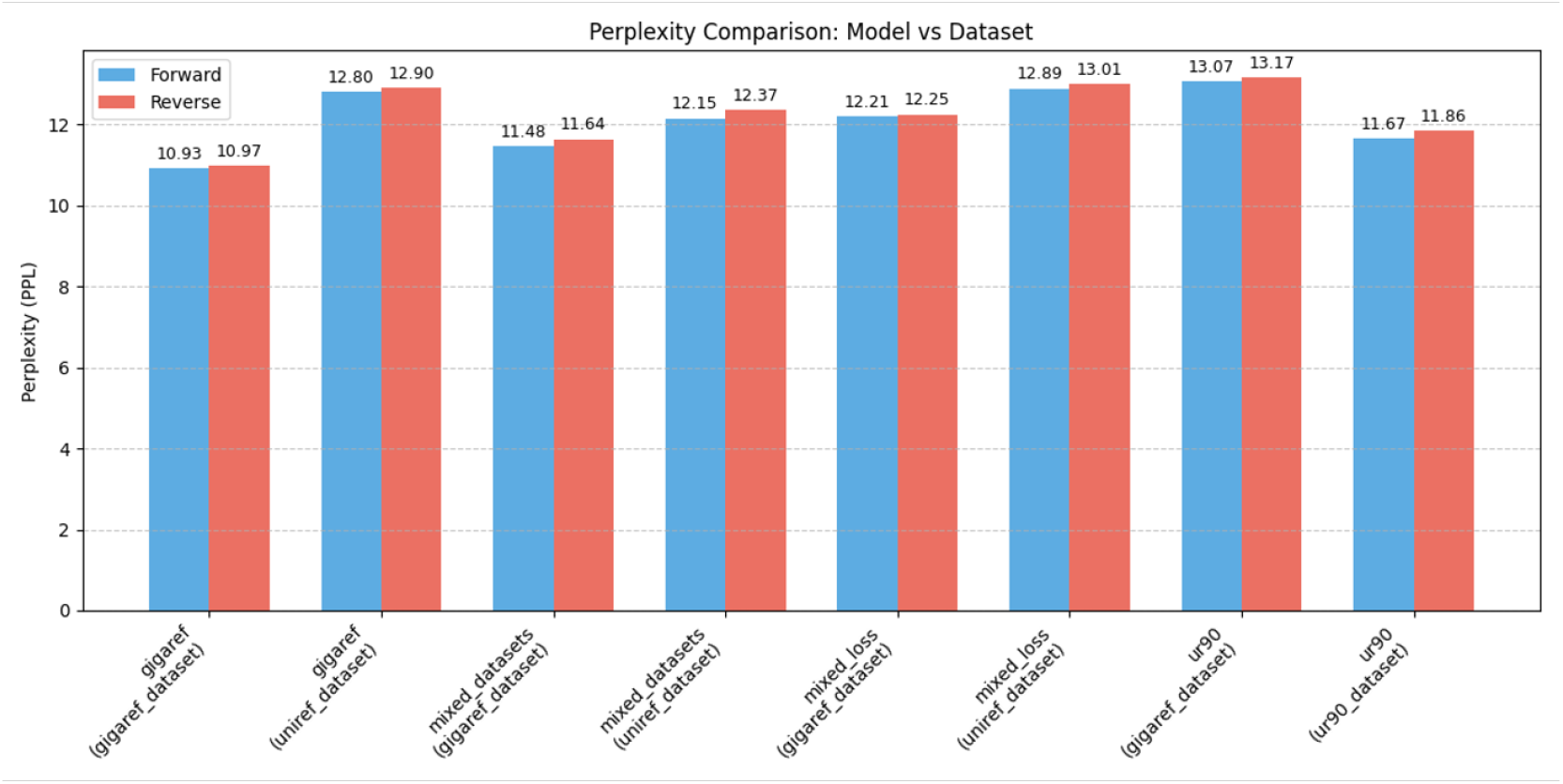
Validation perplexity on Gigaref and Uniref90 validation dataset across the four 112M-parameter models trained on: (1) GigaRef sequences only, (2) 50/50 mixed batches, (3) GigaRef with a weighted loss to up weight Uniref sequences, and (4) Uniref90 (ur90) sequences only.

**Figure 6:**
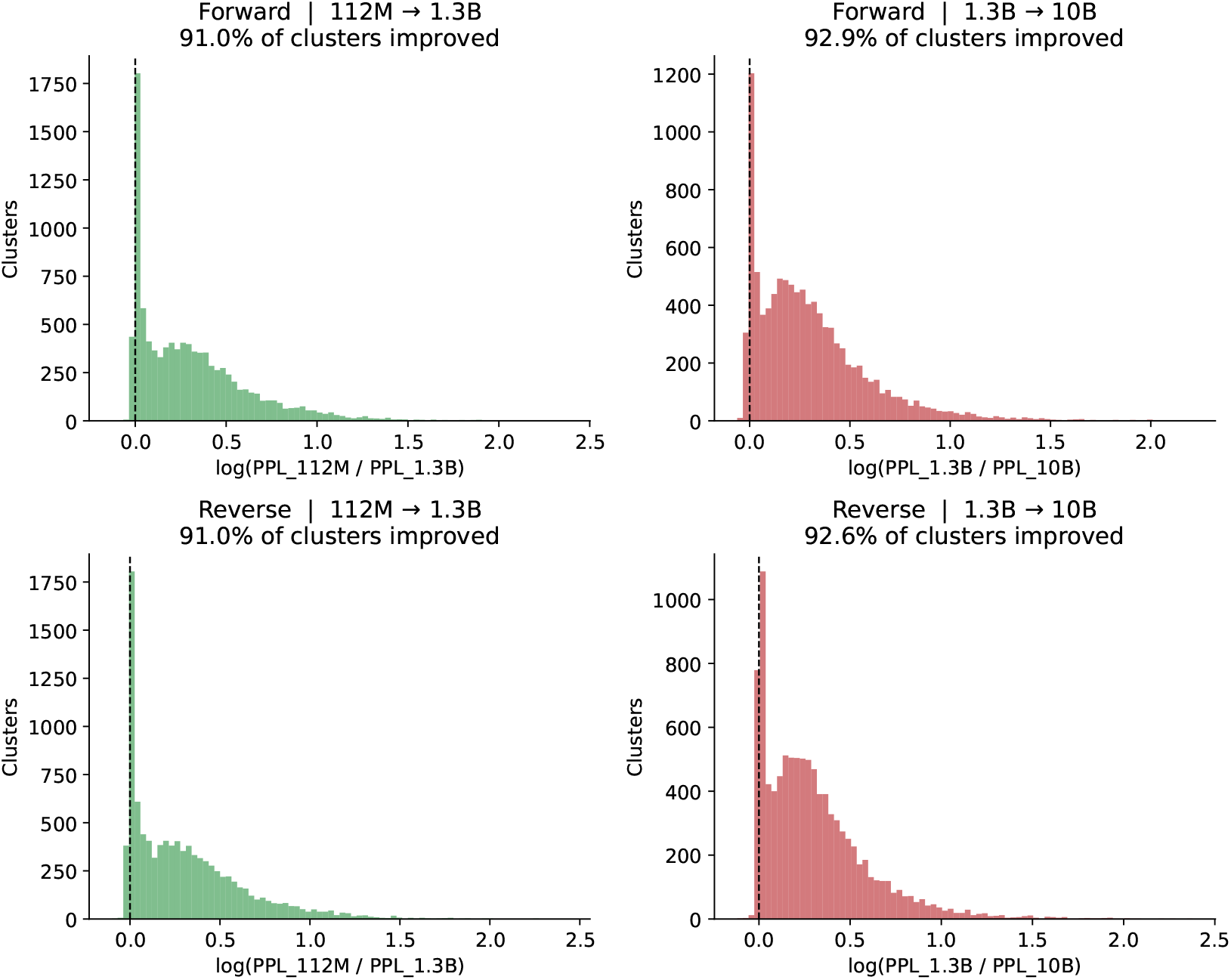
Per-cluster perplexity improvement across model scales. Log-ratio of per-cluster perplexity between adjacent model sizes, 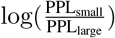, for forward (N-to-C) (top) and reverse (C-to-N) terminal (bottom) direction. Values greater than zero indicate clusters where the larger model assigns lower perplexity. Both scaling steps improve perplexity on over 91% of clusters, with the 1.3B→10B transition showing a broader distribution of gains, reflecting more uniform improvement across the sequence space.

### 3.4 Few-shot prompting enables structurally consistent and fitness-aware protein generation

We first investigate the capacity of ProtGPT3-MSA to generate structurally consistent proteins when conditioned on few-shot prompts composed of homologous sequences drawn from clusters spanning a range of sequence identity levels (30–100%). We observe high structural consistency (TM-scores > 0.5) across all sequence identity ranges (Fig. 7b in Appendix), suggesting that ProtGPT3-MSA can generate structurally consistent homologs even when conditioned on prompts spanning a broad range of sequence identities. Next, we benchmark ProtGPT3-MSA against ProteinGym [33] (see Table 2 in Appendix). In line with previous works [58, 51], we find that training with homologous sequences improves predictive power, despite the substantially smaller number of trainable parameters (e.g., 112M vs 10B). Interestingly, further enriching the input with information from additional homologs through an ensemble strategy, improves performance, directly suggesting that evolutionary context can substantially enhance fitness estimation given the same parameter budget, resulting in the best performing model in the ProtGPT family for such task (see Appendix A.4 for further details).

**Figure 7:**
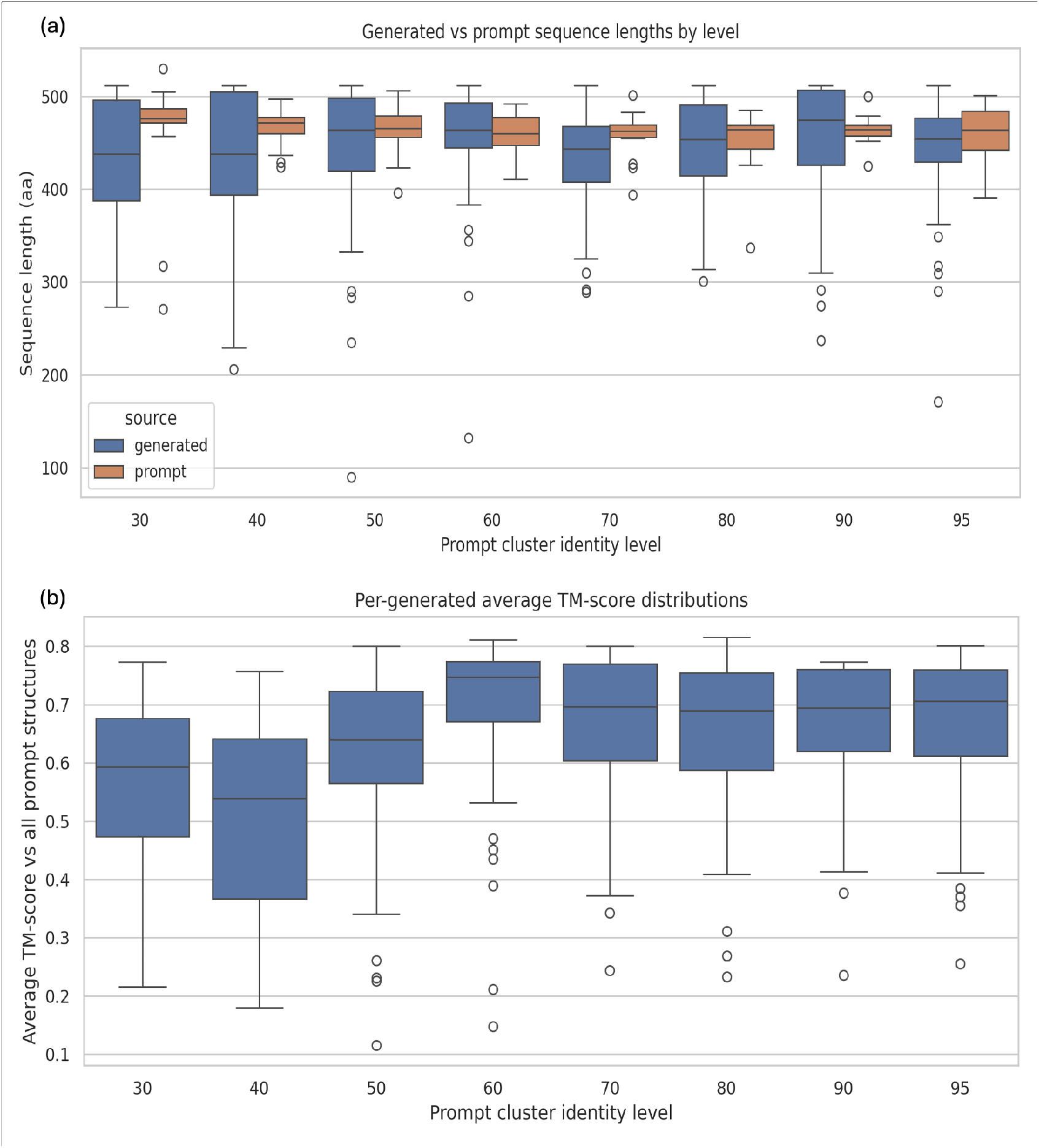
a) Few-shot prompting of ProtGPT3-MSA with homologs spanning 30–100% sequence identity produces proteins that remain within the expected sequence length as the prompt across the full identity range. b) Few-shot prompting of ProtGPT3-MSA with homologs spanning 30–100% sequence identity produces proteins that remain structurally consistent with the prompt across the full identity range.

We next ask whether few-shot prompting can steer ProtGPT3-MSA toward variants with specific properties. Specifically, we attempt to steer generation toward proteins with higher measured stability. Using ten DMS libraries from Tsuboyama et al. [52], we construct prompts containing *K* = 15 variants sampled from specified stability percentiles and generate one additional sequence per prompt.

We score only “hits”, i.e., generated sequences that appear in the corresponding DMS library but were not included in the prompt, allowing us to assign an experimental stability measurement to each generation. Across 12,000 generations, the model produces hits most frequently at *T* = 0.8 (~ 28% hit rate), while higher-temperature sampling reduces recovery (~14% at *T* = 1.0; Fig. 9). Among recovered hits, generated variants are substantially more stable than the prompt mean, with an average gain of Δ = +0.479 DMS units (*t* = 27, *n* = 2,744 hits; Fig. 8–9). This upward shift is visible across prompt percentiles and sampling temperatures, with variability depending on the prompting strategy (tight, linear or concave, see Fig.1e), but irrespective of the order of the sequences (sequences ordered by increasing stability do not lead to generations of higher stability (Fig. 10)). Together, these results suggest that ProtGPT3-MSA can use few-shot prompts to bias generation toward user-defined functional regions of the local sequence space.

**Figure 8:**
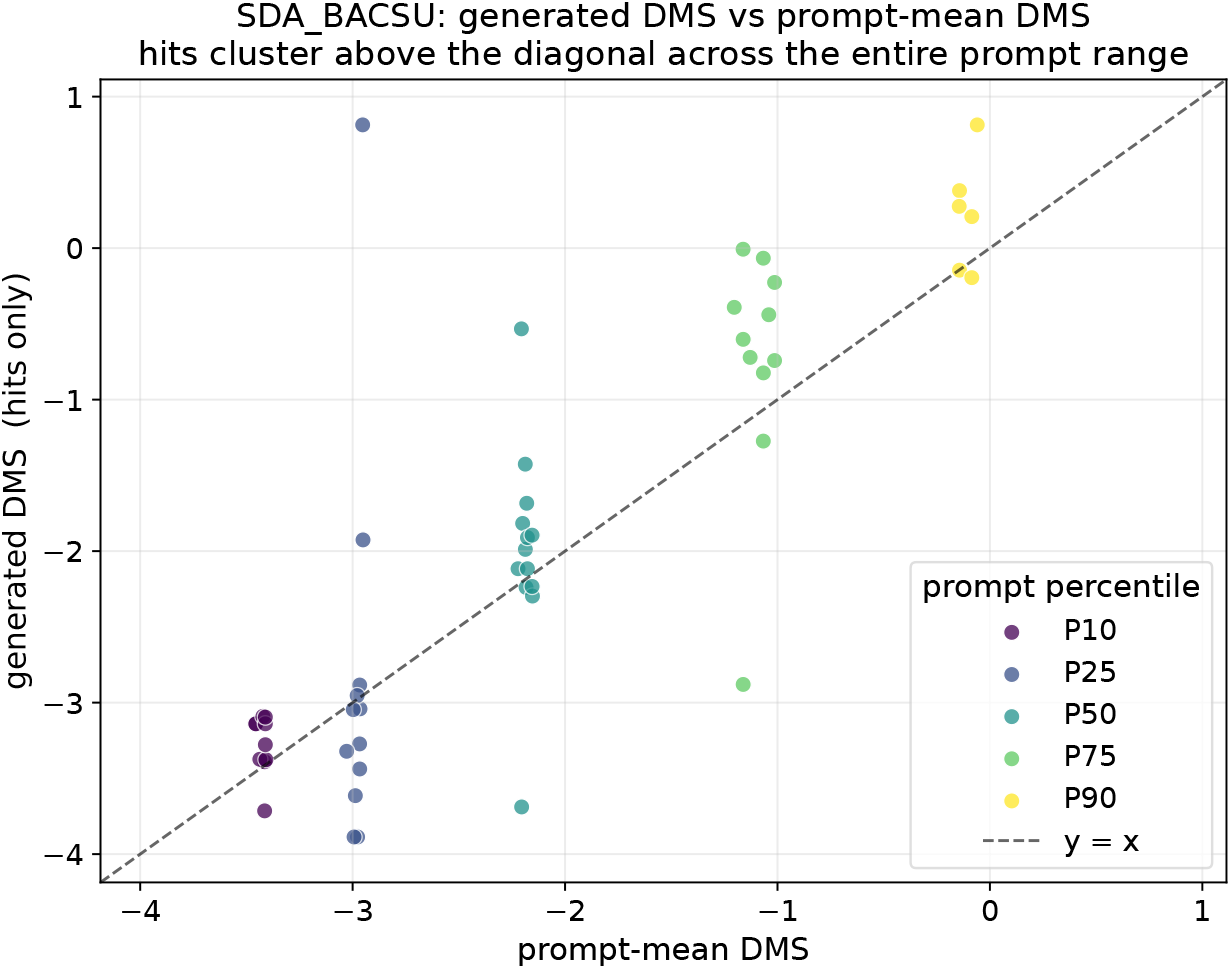
SDA_BACSU illustrative example: generated DMS score versus prompt-mean DMS score for hits only. Points above the diagonal indicate generations with higher measured stability than the prompt mean.

**Figure 9:**
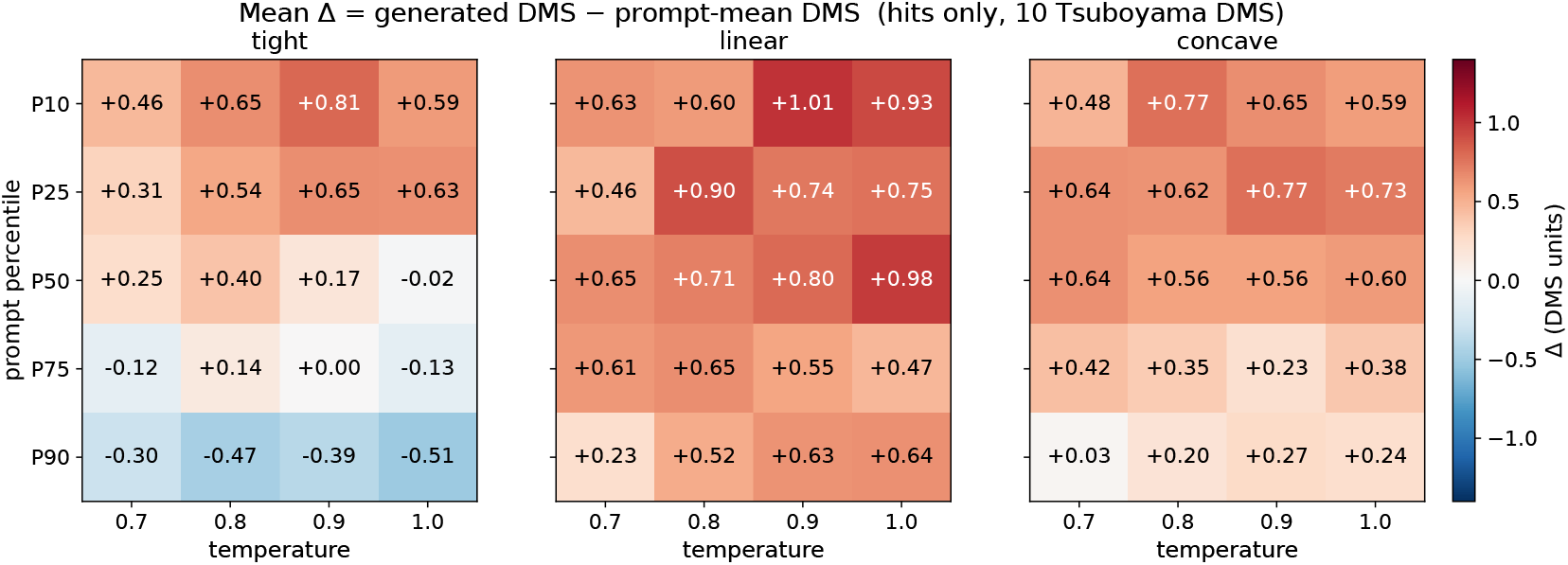
Mean generated-minus-prompt-mean DMS score, computed over hits, across prompt percentile, strategy, and temperature. The linear strategy maintains positive gains across percentiles, whereas tight high-percentile prompts saturate and become negative at P90.

**Figure 10:**
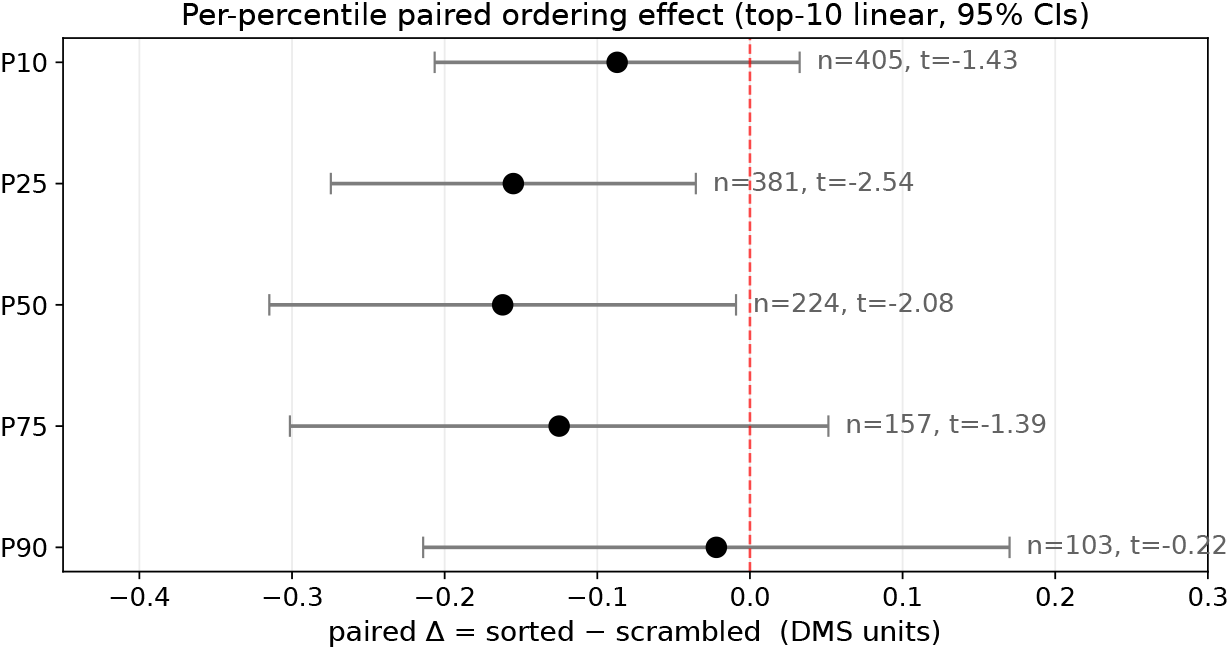
Paired sorted-versus-scrambled prompt control by prompt percentile. Values denote the paired difference in generated DMS score between sorted and scrambled prompts. Negative values indicate that sorted prompts perform worse than scrambled prompts.

**Figure 11:**
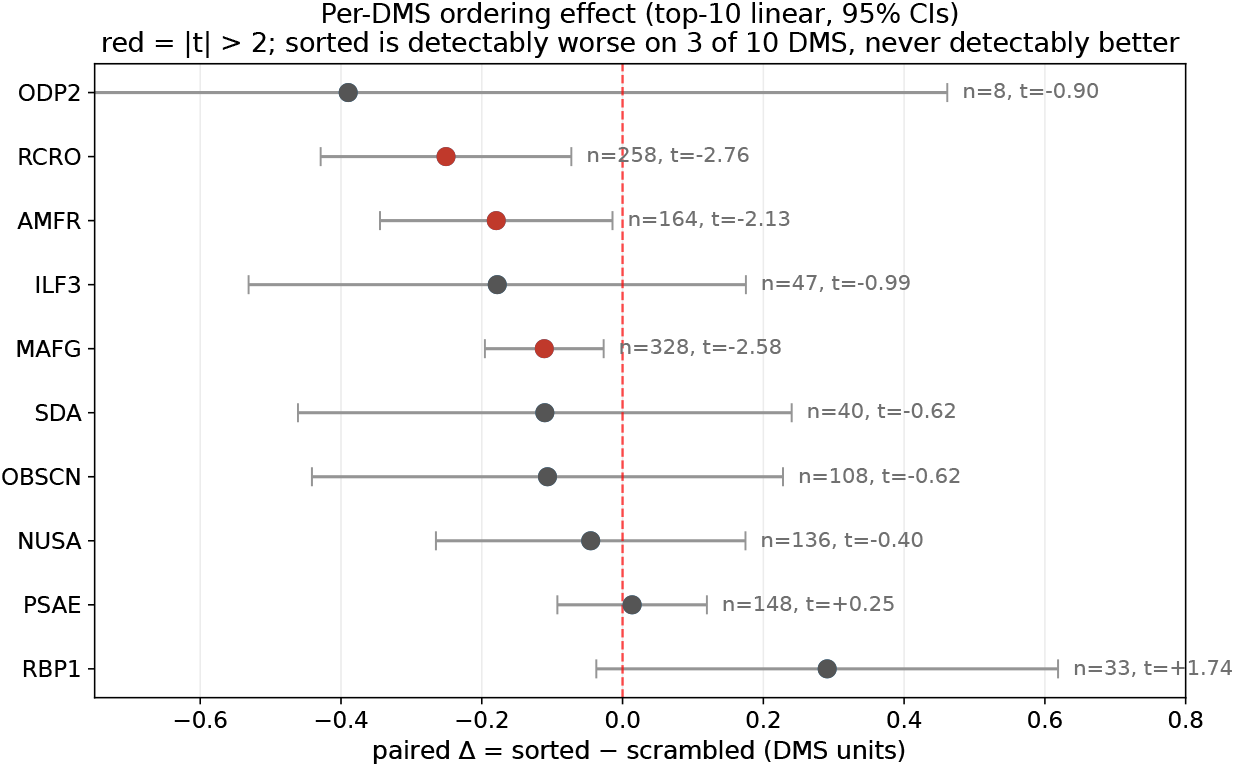
Paired sorted-versus-scrambled prompt control by DMS library. Sorting prompt sequences by DMS score is detectably worse for several libraries and is never detectably better.

## 4 Prompt-based vs SFT for conditional protein generation

We investigate whether ProtGPT3-MSA enables an alternative paradigm for conditional protein generation that does not require task-specific fine-tuning. In contrast to traditional single-sequence autoregressive models, which are often fine-tuned (SFT) on a target protein family to generate family-consistent sequences, MSA models can instead be prompted with homologous sequences from the target protein family to induce conditional generation [e.g., 60, 58, 51]. Despite this conceptual advantage, a systematic comparison between these two approaches remains underexplored. To address this, we design a multi-step evaluation pipeline that compares (i) SFT single-sequence models and (ii) few-shot prompted MSA models for conditional protein generation across diverse protein families. For each tested protein family, we construct train/validation splits and evaluate generated sequences using complementary metrics capturing sequence-, structure-, and family-level consistency: maximum sequence identity to held-out validation sequences, maximum structural similarity (TM-score) to held-out validation sequences as well as family-wide MSA statistics through HHM profile–profile comparisons and per-column KL-divergence (see Appendix A.8 for an in-depth description of each of these comparisons). For a fair comparison with SFT of single-sequence models, all sequences were passed to ProtGPT3-MSA in the *unaligned* mode across all comparisons (i.e., single-sequence models were not trained to process aligned sequences), although the aligned mode showed comparable performance (Appendix Fig 17). We apply this evaluation pipeline across multiple scenarios: (i) 10 randomly sampled MSAs of varying depth (100–1000 sequences) drawn from held-out validation data unseen during ProtGPT3-MSA pre-training, (ii) 10 randomly sampled MSAs from structurally curated datasets disjoint from ProtGPT3-MSA training set i.e, OpenProteinSet/PDB [1] (in Appendix A.10) iii) two targeted case studies involving enzyme families, cutinases (in Appendix A.13) and defluorinases, including experimental validation for the latter.

While SFT can leverage as many homologous sequences from a target protein family as available, ProtGPT3-MSA is limited to 15 homologous sequences at inference time. To overcome this limitation, we implement *prompt ensembling*: *M* independent sets of 15 homologs are sampled as prompts from the target family, the model scores each in parallel, and the resulting next-token distributions are merged into a single sampling distribution at each decoding step. We evaluate three different ensembling approaches (Appendix A.7), with “Max-Ensemble” performing best, though only marginally better than randomly sampling 15 homologs. Max-Ensemble scores each prompt of 15 homologous sequences *ℝ*^(*m*)^ by the mean per-token log-probability the model assigns to it:

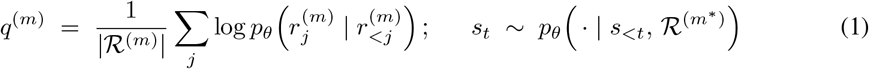

intuitively measuring how well each context set aligns with the model’s learned sequence distribution. Next, the single highest-scoring set 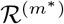, where *m*^*\**^ = arg max_*m*_ *q*^(*m*)^, is used as the prompt to condition the next token prediction at each generation step *t*. While the strong performance of random prompt sampling alone is consistent with prior work [58, 51], Max-Ensemble demonstrates that exploiting model confidence to select higher-quality context sets yields a small but consistent improvement in conditional generation, making it our default prompting strategy for all comparisons.

### Conditional generation comparison for held-out validation MSAs

As a first comparison, we evaluated SFT versus prompting ProtGPT3-MSA across 10 protein families not seen during ProtGPT3-MSA pretraining. To this end, we sampled 10 held-out MSAs from ProtGPT3-MSA pretraining validation set, using a stratified, log-spaced scheme to ensure coverage across MSA depths (100–1000 sequences) and a range of sequence (30% − 60%) and structural (20% − 70%) diversity (see Appendix A.9). Each MSA was treated as an independent evaluation instance. For each sampled MSA, we constructed a train/validation split, and used the training split both to fine-tune ProtGPT3-112M and to prompt ProtGPT3-MSA. We then generated 90 sequences per model (10 sequences for each combination of top-p ∈ {0.7, 0.9, 1.0} and temperature ∈ {0.8, 0.9, 1.0}). Finally, for each MSA, we compared the generated sequences from both models against the corresponding held-out validation split (i.e., sequences not used for either SFT or prompting) using metrics of sequence-, structure-, and family-level consistency (Appendix A.8). Fig 2a-f shows prompting ProtGPT3-MSA outperforms SFT of ProtGPT3-112M by generating sequences that more closely reflect the target protein family across all measured metrics, despite ProtGPT3-MSA not having been trained on those families. Here, we compared ProtGPT3-MSA vs SFT ProtGPT3-112M to ensure an equal number of parameters across the two models. In Fig 15 in the Appendix, we show the exact same comparison for ProtGPT3-MSA vs SFT ProtGPT3-10B, showing ProtGPT3-MSA still performs competitively to much larger models, especially on the family-level consistency metrics. To get a concrete measure of how well ProtGPT3-MSA does on the family-level consistency metrics (i.e., HHalign and positional KL-divergence), we included two baselines, “Bootstrap MSA” and “Scrambled MSA”, respectively approximating upper and lower bound performance on these metrics. Specifically, Bootstrap MSA samples 30% of each true MSA sequences, performing the analysis for sequences that come from the true family distribution for each MSA. Conversely, “Scrambled MSA” provides a negative control, by running the family-level consistency analysis for ProtGPT3-MSA generated sequences for each MSA against a different MSA. This ensures the generated sequences by ProtGPT3-MSA do not just match per-column statistics for any MSA. Fig 2d-f shows ProtGPT3-MSA generated sequences achieve family-level consistency metrics close to sequences truly sampled from those family across most tested MSAs (i.e., Bootstrap MSA). In Appendix A.10, we show similar performance of ProtGPT3-MSA for 10 MSAs randomly sampled from the Protein Data Bank subset of OpenProteinSet [1].

### Conditional generation comparison for Low-N design of defluorinase enzymes

Defluorinases were selected as a test case for experimental characterization under extreme data scarcity. These enzymes cleave carbon–fluorine bonds found in persistent pollutants such as PFAS. Defluorinases occur in the (S)-2-haloacid dehalogenase and α/β-hydrolase superfamilies, where defluorinating activity evolved independently [56]. Each superfamily has less than 10 experimentally annotated defluorinases, that are also highly similar to dechlorinases - enzymes cleaving carbon-chloride bonds. Because defluorinases are highly similar to dechlorinases and their distinguishing molecular features remain unclear [36], family-level datasets remain too ambiguous, whereas experimentally confirmed defluorinases are too scarce. Therefore, we set out to compare the generation of defluorinases from the (S)-2-haloacid dehalogenase superfamily via supervised fine-tuning of the single-sequence models and few-shot prompting of ProtGPT3-MSA with the 7 defluorinases identified and characterized by Probst et al. [36]. The two approaches were evaluated computationally and experimentally. Few-shot generation with ProtGPT3-MSA achieved 137-fold and 68.5-fold higher computational success rates (Appendix A.12) than fine-tuned ProtGPT3-1.3B and ProtGPT3-10B, respectively, while ProtGPT3-112M did not generate any successful sequence (Fig. 2g). ProtGPT3-MSA performed consistently better on average than the fine-tuned single sequence models, which produced successful sequences only as outliers from the distribution of their generations (Fig 18 in Appendix A.12). We observed similar behaviour for the targeted generation of cutinases (see Appendix A.13). For experimental validation, we selected all sequences passing the filters for the single-sequence models, providing 3 and 7 sequences for the 1.3B and 10B models. As ProtGPT3-MSA had many more sequences that passed the initial selection criteria than what we could test experimentally, we applied further filtering steps (see Appendix A.12) and chose the 10 highest-ranking sequences for experimental testing. After cloning and purification, all 18 out of the 20 successfully cloned sequences showed protein yields after expression and purification detectable by Bradford assay, indicating the successful soluble, expressible candidate defluorinases (Fig. 2i & Fig. 19).

## 5 Inference-time compute strategies in promptable pLMs

Auto-regressive multiple-sequence protein language models, such as ProtGPT3-MSA, provide a natural setting for exploring inference-time compute strategies compared with single-sequence models. Because generation is organized over sets of homologous sequences rather than individual amino acids, the model induces a structured, sequence-level search space that is well suited to trajectory-based selection. This structure is particularly advantageous because many objectives in protein design are defined at the level of full sequences, allowing scoring functions to be directly incorporated into the sampling procedure. To exploit this structure, we adapt a Feynman-Kac-style sequential Monte Carlo (FK-SMC) procedure to sample sequences with a target objective from ProtGPT3-MSA (see Algorithm 1 in Appendix). Given an initial prompt consisting of one or more concatenated homolog sequences, we let ProtGPT3-MSA branch N candidate trajectories, where each trajectory is a multi-step rollout over K homolog sequences generated autoregressively. We evaluate each trajectory using a sequence-level scoring function applied over the full K-step continuation, retain the first homolog sequence from the highest-scoring trajectory, append it to the prompt, and repeat. Importantly, because ProtGPT3-MSA was trained to generate sequences evolutionary related to its prompt, the search is implicitly constrained to the neighborhood of the starting sequence (i.e., generated sequences are biased towards biologically plausible homologs rather than arbitrary sequences). The key assumption is that if variants with the desired property exist within this homolog space, the sequential FK Monte Carlo procedure can steer generation toward them.

We validate the FK-SMC procedure in a controlled setting by steering generation between pairs of homologous sequences with moderate structural similarity (TM-score ∈ [0.5, 0.6]), using ESM2 [27] cosine similarity to a designated target as the scoring function (details in Appendix A.14). Across 10 randomly sampled pairs, cosine similarity to the target increases consistently at each step (Fig. 3a), confirming that the procedure reliably steers generation within the model’s learned homolog space. The UMAP trajectories further illustrate this progressive movement toward the target in embedding space (Fig. 3b). While ESM2 cosine similarity and TM-score are not expected to correlate directly [35], several pairs nonetheless show a modest increase in TM-score (Fig. 21 in Appendix), suggesting that embedding-space steering can carry some structural signal.

## 6 Aligning single-sequence pLMs for improved sequence generation quality

A common failure mode of LLMs is repetitive generation, which in single-sequence protein language models often manifests as low-complexity sequences [57, 18, 4, 10]. Although scaling partially alleviates this issue (Fig. 1d), all models still generate an excessive percentage of low-complexity sequences. To reduce this bias, we implement a reinforcement-learning-based alignment pipeline for each single-sequence model. We first generate 80,000 sequences across combinations of top-*p* ∈ 0.6, 0.8, 0.9, 1.0 and temperature ∈ 0.6, 0.8, 0.9, 1.0 from each model, and score each sequence based on both the fraction of low-complexity regions and pLDDT predicted by ESMFold [27]. Sequences with pLDDT *>* 0.7 and fewer than 25% low-complexity residues are treated as positive examples, with all others treated as negatives. The pLDDT criterion acts as a proxy for structural plausibility, preventing trivial optimization toward random high-complexity amino-acid strings. We then apply Direct Preference Optimization (DPO) [38] to bias generation toward positive sequences, repeating the procedure for up to three iterations (see Appendix A.15 for details). Across all models, alignment substantially improves generation quality, increasing the proportion of high-complexity sequences (Fig. 3c) while maintaining high pLDDT scores (Fig. 22b in Appendix). In particular, the fraction of low-complexity generations decreases by more than 20% for the 112M and 1.3B models, and by nearly 50% for the 10B model. Alignment also increases sequence diversity across all scales (Fig. 3d), while pre-training validation perplexity remains largely unchanged (Fig. 22c in Appendix), suggesting minimal catastrophic forgetting of the pretraining sequences. Notably, a single iteration of the alignment pipeline reduces low-complexity generations below 5% for ProtGPT3-10B, suggesting that larger models are particularly responsive to alignment and may be easier to steer toward biologically realistic sequence distributions [e.g., 4].

## 7 Conclusion and limitations

We introduce ProtGPT3, an open-source family of autoregressive protein language models spanning 112M to 10B parameters, together with ProtGPT3-MSA, a 112M-parameter model for homolog-conditioned generation. All models are integrated with the HuggingFace ecosystem and released with efficient training and generation code, lowering the barrier to customizable protein language models. Across scale, larger single-sequence models improve unconditional generation, diversity, and variant-effect prediction. We also show that post-training alignment can reduce low-complexity generation while preserving sequence diversity, suggesting that, similarly to NLP models, pLMs benefit from large-scale alignment procedures to mitigate broad sampling biases beyond task-specific optimization. A central finding is that homolog-conditioned prompting can replace task-specific fine-tuning in many conditional generation settings. While still requiring homolog sequences, ProtGPT3-MSA reduces the burden of task-specific fine-tuning, as only a small set of related sequences is needed to generate family-consistent proteins without updating model weights. This performs competitively with, and often outperforms, supervised fine-tuning of much larger single-sequence models. Experimentally, ProtGPT3-MSA achieved substantially higher computational success rates than fine-tuned baselines and produced designs that were expressed and soluble under our assay conditions. Our study has several natural limitations, most of which point to direct next steps. Although soluble expression is a necessary prerequisite for enzyme activity and is itself a challenging design objective, enzymatic defluorination activity remains to be tested experimentally. Similarly, the FK-SMC experiment demonstrates controllable embedding-space steering, but its utility for optimizing real biological objectives remains to be established in future case studies. A broader open challenge, shared by both inference-time steering and RL-based alignment, is the identification of reliable computational proxies for experimentally relevant protein properties. Despite these limitations, our results suggest that prompting, alignment, and inference-time compute represent a promising framework for controllable protein generation. We hope ProtGPT3 and ProtGPT3-MSA provide a useful open foundation for future work on scalable and controllable protein design.

## 8 Acknowledgements

The authors thank the Barcelona Supercomputing Center for GPU resources provided through projects EHPC-DEV-2025D11-089, EHPC-DEV-2025D10-074, BCV-2025-3-0055, and BCV-2025-2-0034. The authors acknowledge the scientific support and HPC resources provided by the Erlangen National High-Performance Computing Center (NHR@FAU) of the Friedrich–Alexander-Universität Erlangen–Nürnberg (FAU) under the NHR project b114cb (UID 210235).

MG acknowledges funding from project CPP2023-010699, funded by MCIU/AEI/10.13039/501100011033 / FEDER, UE. FS is supported by a fellowship from the “la Caixa” Foundation (ID 100010434, fellowship code DFI25-00620F). LM acknowledges support from the predoctoral program AGAUR-FI ajuts (2025 FI-3 00065) Joan Oró, of the Department of Research and Universities of the Generalitat of Catalonia, as well as the European Social Plus Fund. NF acknowledges support from a Ramón y Cajal contract RYC2021-034367-I, funded by MCIN/AEI/10.13039/501100011033 and by the European Union NextGenerationEU/PRTR. We acknowledge the support of the Spanish Ministry of Science and Innovation through the Centro de Excelencia Severo Ochoa (CEX2020-001049-S, MCIN/AEI/10.13039/501100011033), and the Generalitat de Catalunya through the CERCA program. We gratefully acknowledge support from the project “Proyectos de Generación de Conocimiento” PID2022-139006NA-I00 funded by MCIU/AEI/10.13039/501100011033 / FEDER, UE and the BBVA Foundation Grants for Scientífic Research Projects 2023 through its Leonardo Creators 2023 grant (ID LEO23-2-10299-CCD-CIA-15). We acknowledge support from the the European Union’s Horizon Europe under the grant agreement No 101165231.

Views and opinions expressed are, however, those of the author(s) only and do not necessarily reflect those of the European Union. Neither the European Union nor the granting authority can be held responsible for them.

## A Technical appendices and supplementary material

### A.1 Model architecture: Sparse vs Dense

Here we report the training and inference comparison between a 112M-parameter Mixtral-style (sparse) mixture-of-experts (MoE) model [20] and an equivalent 112M-parameter llama3-style (dense) model [15]. Both models were trained on the exact same infrastructure (4 x H100 NVIDIA GPUS) on 9.8B tokens following optimal scaling law for AR pLMs from Cheng et al. [10]. Fig. 4a,b shows the (sparse) Mixtral architecture achieves marginally worse validation perplexity, PPL (i.e, both forward and reverse), while enabling faster training (≈1*/*3 faster) relative to the (dense) LLama3 architecture. The validation perplexity was based on a held-out set of protein sequences, built such that all sequences were at least 50% different from the training sequences in terms of sequence identity (0.8 coverage). Additionally, inference of the validation sequences is ≈ 1*/*2 faster in the (sparse) Mixtral architecture compared to the LLama3 one (see Fig. 4c) based on the same infrastructure (a single H100 NVIDIA GPU).

### A.2 Pretraining sequences

All models were trained with publicly available datasets. In table 1, we report the number of protein sequences used to train each single-sequence model across the two employed datasets, 1) Uniref90 [53] and 2) Gigaref [60]. The number of sequences used for each model was based on optimal scaling laws for sparse protein language models [4], enforcing 50% of tokens from each dataset for the 112M and 1.3B parameters models and, 70% Gigaref with 30% Uniref90 tokens for the 10B parameter model. During training each sequence was randomly assigned a generation direction (forward) N-to-C or (reverse) C-to-N. The special tokens 1 and 2 were prepended to each sequence to respectively denote the N-to-C and C-to-N directions.

**Table 1:** Number of protein sequences used to train each single-sequence model across the two employed datasets.

| Model size | Uniref90 | Gigaref | N. tokens |
| --- | --- | --- | --- |
| 112M | 15M | 28M | 9.8B |
| 1.3B | 64M | 120M | 43B |
| 10B | 174M | 643M | 170B |

**Table 2:**
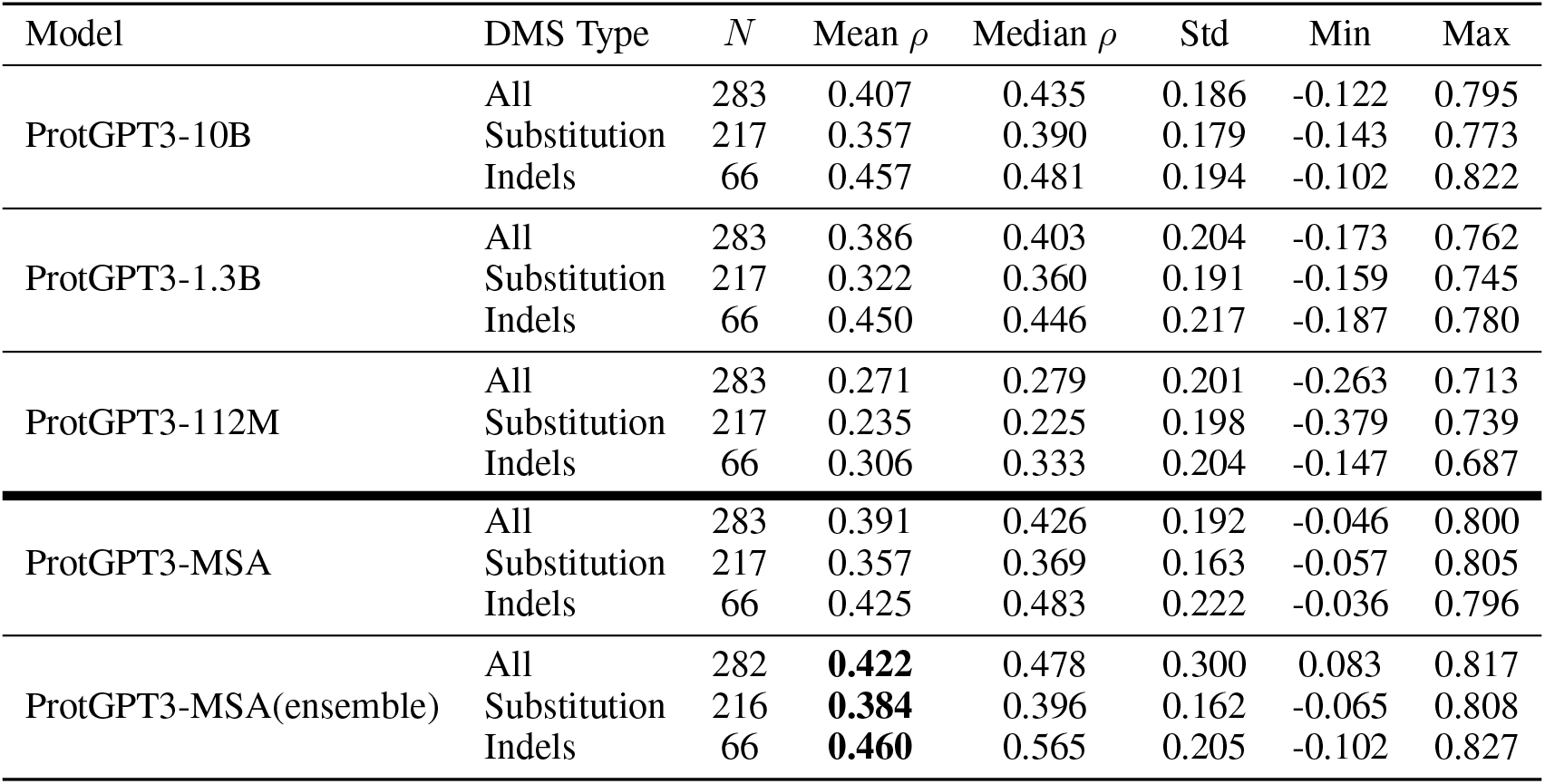
Spearman correlation (*ρ*) between model log-likelihoods and experimental fitness scores on the ProteinGym DMS benchmark.

The MSA model was using the OpenProteinSet Uniclust30 dataset [1]. The 16M Uniclust30 MSAs were filtered out for having less than 16 sequences and for any containing sequence having more than 1024 amino acids. This resulted in approximately 8.5M training MSAs, from each a set 16 sequences were sampled without replacement and this process was repeated 15 times, always randomizing the order in which the 16 sequences were concatenated. In total, this led to training ProtGPT3-MSA on approximately 560B tokens.

### A.3 Validation per-cluster perplexity analysis

To assess how uniformly model quality improves with scale, we evaluated each model perplexity at the level of 50% sequence clusters rather than computing a single aggregate perplexity across the entire validation set. We ran this analysis for held-out validation clusters, those sequences had at most 50% sequence identity to the training set of each model. Concretely, for each cluster in the held-out validation set we computed the mean token-level negative log-likelihood across all member sequences, evaluated for both forward (N-to-C) and reverse (C-to-N) terminal direction, and converted this to a per-cluster perplexity. This cluster-level view allows us to distinguish between a model that improves uniformly across the sequence space and one that improves only on clusters it already models well, a distinction that is invisible when reporting a single validation average.

Fig. 6 shows the log-ratio of per-cluster perplexity (PPL) between adjacent model sizes, 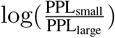 for both forward (N-to-C) and reverse (C-to-N) terminal direction. Values to the right of zero indicate clusters where the larger model assigns lower perplexity, i.e. models the sequences better. Both transitions, 112M to 1.3B and 1.3B to 10B, show strongly right-skewed distributions, with over 91% of clusters improving at each step and in both orientations. Notably, the 1.3B→10B histograms display a broader, more uniformly spread distribution compared to the sharper spike near zero seen for 112M→1.3B, indicating that the gain from the largest model is more evenly distributed across clusters rather than concentrated on a small subset. The near-identical results for forward and reverse orientations confirm that the bidirectional training objective is learning consistent representations regardless of sequence direction.

### A.4 ProteinGym benchmarking

In Table 2, we report the results of benchmarking each model on ProteinGym [33]. The mean Spearman correlation was computed using the official ProteinGym script, `performance_DMS_benchmarks.py`. The median, standard deviation, minimum, and maximum were computed over the DMS-level Spearman correlations. Assemblies were constructed from sets of 16 orthologous sequences randomly sampled from the precomputed MSAs provided by ProteinGym. Each sequence was evaluated in both the forward and reverse directions using the corresponding direction tokens. The resulting log-likelihoods were averaged across directions. The change of fitness determined by the mutation (Δ*f*) was computed as a paired log-likelihood difference between each mutant sequence and its corresponding wild-type sequence. For each replicate, the mutant and wild-type sequences were each inserted as the 16th sequence and scored using the same set of 15 orthologous context sequences, ensuring a paired comparison. For each DMS assay, we evaluated a random subsample of up to 1,000 variants. The final fitness score was obtained by averaging the paired differences over 10 replicates.

For each variant, we used a paired scoring procedure in which the mutant and wild-type sequences were evaluated using the same sampled assembly context. Let *d* ∈ {fwd, rev} denote the sequence direction. We first computed the assembly-averaged sequence score

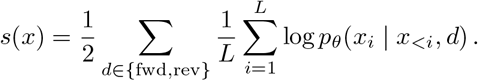

In case of ProtGPT3-MSA, we have also averaged over 10 different samples of orthologous sequences. The variant fitness score was then defined as the paired difference between the mutant and wild-type scores:

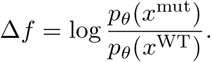

### A.5 Few-shot prompting for functional variant generation

We evaluate whether few-shot prompts can steer ProtGPT3-MSA toward variants with improved experimentally measured stability. For each of ten DMS libraries from Tsuboyama et al. [52], we construct prompts from variants at a target stability percentile and generate one additional sequence. A generation is counted as a hit if it appears in the corresponding DMS library but is not present in the prompt, which allows us to assign an experimental DMS score to the generated sequence. As an illustrative example, SDA_BACSU generations cluster above the diagonal when comparing generated DMS score to the prompt-mean DMS score, indicating that recovered hits are often more stable than the variants used for conditioning (Fig. 8).

We compare three prompt construction strategies. All strategies sample *K* = 15 prompt sequences and sort them by increasing DMS score. The *tight* strategy samples variants from a narrow percentile bucket around the target percentile, the *linear* strategy samples evenly spaced stability targets between the bottom of the library and the target percentile, and the *concave* strategy uses power-law spacing to concentrate samples near the target percentile. Across 12,000 generations, hit rate peaks at *T* = 0.8 and drops at *T* = 1.0; the three strategies are similar in hit rate across temperatures (Fig. 9). Among hits, generated variants have higher DMS scores than their prompt means on average, with mean Δ = +0.479 DMS units (*t* = 27, *n* = 2,744 hits). This effect is visible both in within-DMS normalized generated scores across prompt percentiles (Fig. 8) and in per-cell generated-minus-prompt-mean DMS differences (Fig. 9). The per-cell analysis reveals substantial strategy heterogeneity: the linear strategy remains positive across the P10–P90 range, while tight prompts become negative at P90, consistent with saturation when the prompt is already drawn from the top of the assayed distribution.

### A.6 Controls for prompt ordering and prompt copying

To test whether the model uses the ordering of prompt examples as an in-context stability trend, we compare sorted prompts to paired scrambled prompts using the same underlying sequences. Sorting does not improve generated stability; instead, sorted prompts are slightly worse on average, and the effect is detectably negative for several DMS libraries and prompt percentiles (Figs. 10– 11). This suggests that the model is not extrapolating from the ascending DMS order of examples. We therefore examine whether generations instead reflect the set of positions mutated in the prompt. Per-position mutation-density heatmaps show that the mutation profile of generated sequences shifts with prompt percentile in several DMS libraries, supporting the interpretation that the prompt acts as a soft positional prior over local sequence variation (Fig. 12). Finally, we quantify verbatim self-copying from the prompt. Self-copy rates among hits are nonzero and vary by strategy and percentile, motivating the exclusion of prompt-contained sequences from the hit definition and providing a quality-control diagnostic for few-shot generation (Fig. 13).

**Figure 12:**
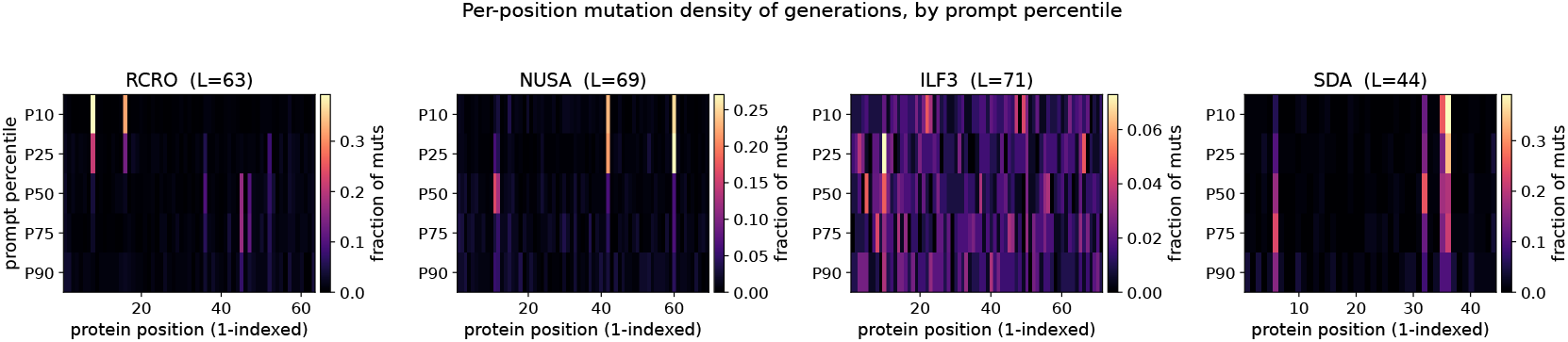
Per-position mutation density of generated sequences by prompt percentile for representative DMS libraries. Generated sequences shift their mutation profiles with prompt percentile, suggesting that the prompt provides a positional prior over tolerated mutations.

**Figure 13:**
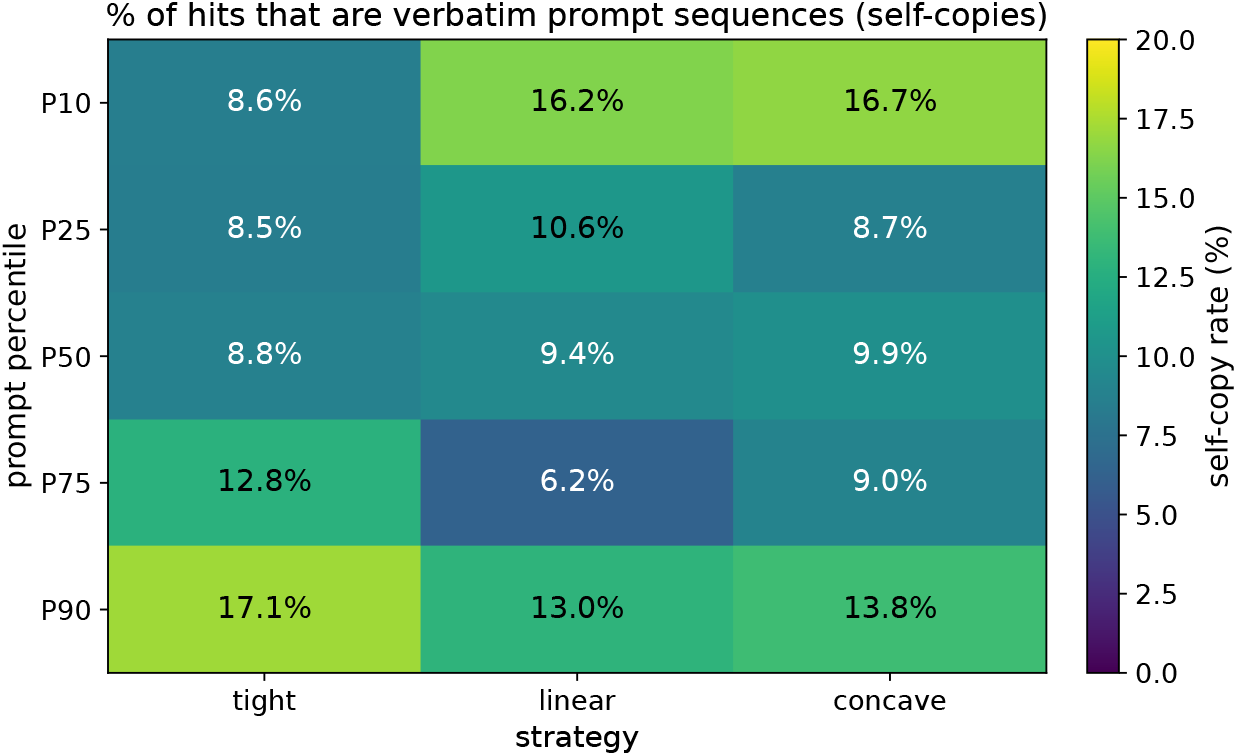
Fraction of hits that exactly copy one of the prompt sequences, stratified by prompt strategy and percentile.

### A.7 Prompt Ensembling Strategies for ProtGPT3-MSA

ProtGPT3-MSA can be prompted with up to 15 homologous sequences (i.e., pretrained on sets of 16 concatenated sequences). However, natural protein families typically contain hundreds to thousands of homologs, far exceeding this context budget. To leverage a larger fraction of the available evolutionary information during conditional generation, we introduce *prompt ensembling*: *M* independent context sets are sampled from the MSA pool (i.e., the target protein family), the model scores each in parallel, and the resulting next-token distributions are combined into a single sampling distribution. We explore three strategies that differ in how this combination is performed, described below. Let *M* denote the number of ensemble members and *V* the vocabulary. At each generation step *t*, every ensemble member *m ∈ {*1, …, *M}* consists of a distinct set of homologous sequences *ℝ*^(*m*)^ sampled from the target protein family (i.e., MSA). Given the current generation step *s*_*<t*_ and context *ℝ*^(*m*)^, the model produces a next-token log-probability vector 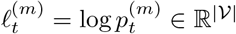, where 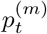 is obtained by applying a softmax with temperature *τ* to the model logits.

**Ensemble** aggregates by averaging in log-space (equivalent to a geometric mean of probabilities) and renormalising:

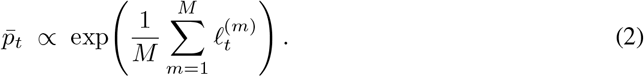

where 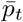 denote the merged (unnormalised) next-token log-probability across the ensemble. The next token is sampled as 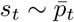 and appended to the shared prefix before the next decoding step. Tokens must be confidently predicted across *all* ensemble members to receive high probability mass, making this the most conservative of the three strategies.

**w-Ensemble** weights each member’s distribution by the model’s own confidence in its context. Specifically, a scalar quality score *q*^(*m*)^ is assigned to each prompt by computing the mean per-token log-probability over the prompt tokens *ℝ*^(*m*)^ in a single forward pass prior to generation:

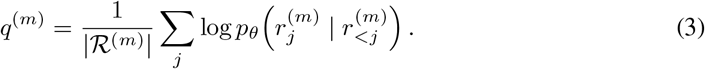

These scores are converted to a weight distribution via softmax, *w*^(*m*)^ = softmax(**q**)_*m*_, and the ensemble distribution is the resulting weighted average in log-space:

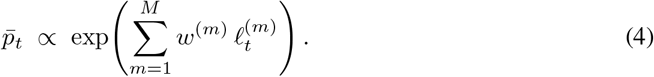

The next token is sampled as 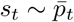 and appended to the shared prefix before the next decoding step. Prompts that the model assigns higher likelihood (i.e., suggesting better alignment with the learned sequence distribution) contribute proportionally more to the final prediction.

**Max-Ensemble** extends w-Ensemble by discarding all but the single most confident ensemble member. Using the same prompt scores *q*^(*m*)^ from Eq. (3), the highest-scoring context set is selected as:

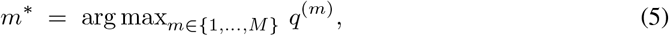

and the next token at each generation step *t* is sampled directly from that prompt’s distribution:

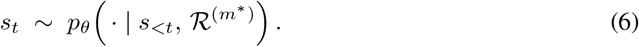

Because all non-selected prompts are pruned before the autoregressive loop begins, Max-Ensemble reduces wall-clock generation time by a factor of *M* relative to running a full *M*-member ensemble, while retaining the benefit of having scored a diverse pool of *M* candidate context sets.

In order to assess which of the 3 prompt ensemble approaches works best, we look at potential difference in the quality of generated sequences across the four approaches. In particular, we assess the quality of generated sequences in terms of MSA per-column statistics, which may better capture fine grained difference across the different approaches compared to more global sequence and structure similarity metrics. For this we randomly sampled 10 MSAs from the Protein Data Bank subset of OpenProteinSet [1]. From each MSA, we randomly sampled 10 sets of 15 homologs (i.e., 150 homologs in total) and used them to (few-shot) prompt ProtGPT3-MSA via one of the 3 ensemble approaches described above, 1) Ensemble, 2) w-Ensemble and 3) Max-Ensemble. Additionally, we also assess the performance of not using an ensemble approach (i.e., No Ensemble) by randomly sampling a single set of 15 homologs and use it to (few-shot) prompt ProtGPT3-MSA. We compared the quality of the generated sequence from the 4 approaches relative to the reference MSA using the positional KL-divergence and HMM profile comparison described in Appendix A.8. Fig. 14 Max-Ensemble performs best, albeit only marginally better than the No Ensemble approach. We find both w-Ensemble and Ensemble perform substantially worse than the two other approaches. We speculate this may be because both w-Ensemble and Ensemble rely on averaging over multiple predictions, which may dilute subtle evolutionary signals detected only by some of the prompts [13].

**Figure 14:**
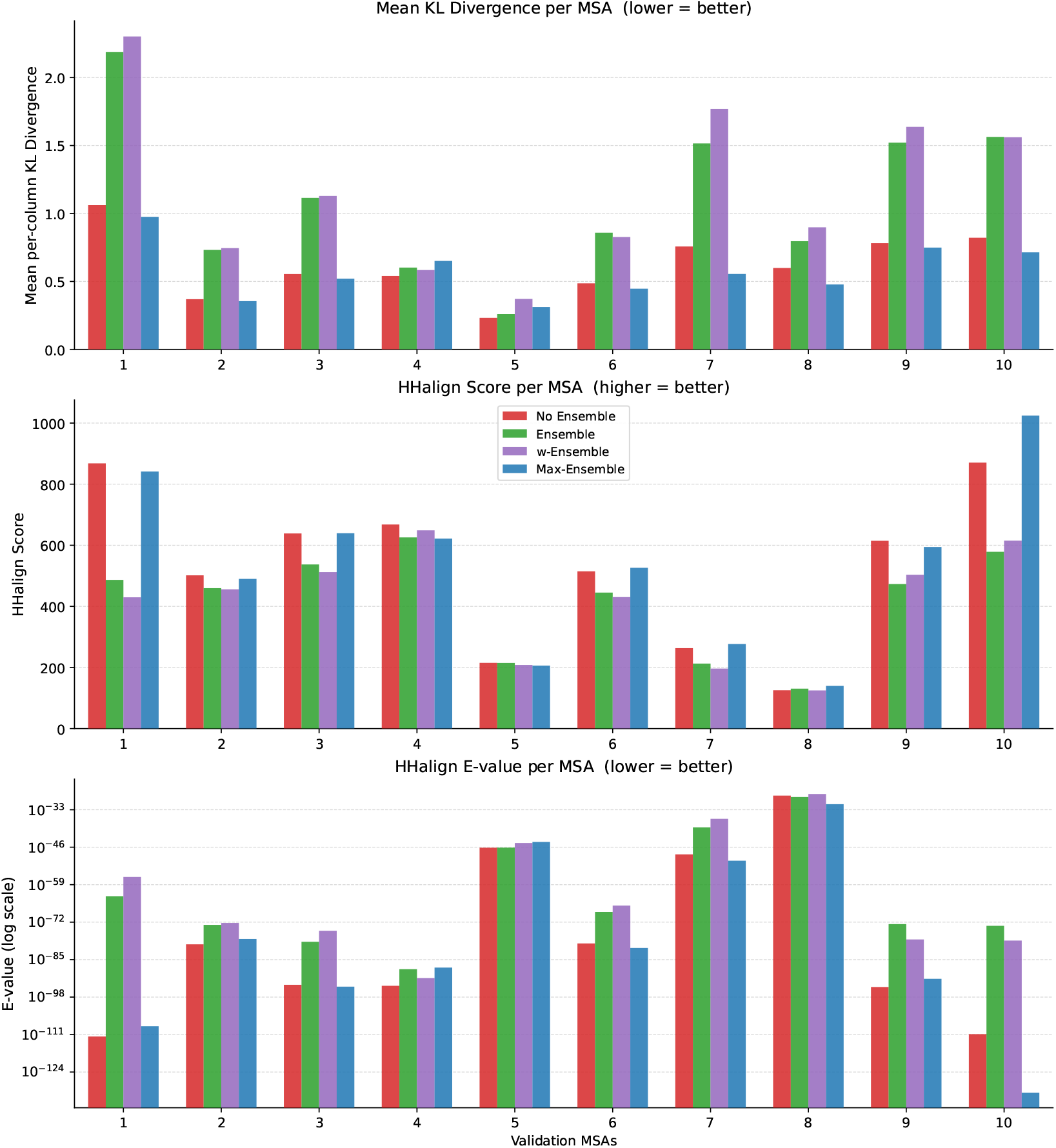
MSA model ensembling approaches comparison for 10 randomly sampled MSAs from the Protein Data Bank subset of OpenProteinSet [1]

### A.8 Evaluation pipeline SFT versus few-shot prompting

We build an evaluation pipeline to compare how few-shot prompting of ProtGPT3-MSA compares to SFT of single-sequence models on target protein family for conditional protein generation. For SFT, each target family is treated as an independent fine-tuning task: we construct per-family train and validation splits, fine-tune the model separately on each, and select the model checkpoint that achieves the lowest validation perplexity for sequence generation. For few-shot prompting of ProtGPT3-MSA, we use Max-Ensemble approach described above to identify the best set of homologs from the target family, which are then used to prompt the model at generation time without any parameter updates. When comparing few-shot prompting of ProtGPT3-MSA to SFT, all ProtGPT3-MSA sequences were generated in *unaligned* modality to ensure a fair comparison (i.e., single-sequence models were not trained to process aligned sequences). However, we include a comparison between *unaligned* and *aligned* generation in Appendix Fig. 17. For both SFT and few-shot prompting, generated sequences are evaluated against the reference family across four complementary metrics (i) maximum sequence identity, (ii) maximum structure similarity, (iii) HMM profile comparison and positional (iv) KL-divergence comparison, each described in detail in the following subsections.

#### A.8.1 Maximum Sequence Identity

To assess the similarity of generated sequences to the reference protein family, we computed the maximum fractional sequence identity between each generated sequence and all reference sequences using MMseqs2 `easy-search` [47]. For each generated sequence, we retained the highest identity match across all reference targets, assigning zero identity to sequences that produced no alignment, so that summary statistics reflect the full generated set. We report the mean maximum sequence identity across all generated sequences within each condition, where lower values indicate greater dissimilarity from known family members.

#### A.8.2 Maximum Structure Similarity

To assess the structural similarity of generated sequences to the reference protein family, we first folded all generated sequences using ESMFold [27] and then computed the maximum TM-score between each predicted structure and all reference structures within the same protein family using Foldseek `easy-search` [54]. For each generated structure, we retained the highest TM-score across all reference targets, normalised by target length, and report the mean maximum TM-score across all generated sequences within each condition. For each predicted structure, we also report ESMFold pLDDT estimates.

#### A.8.3 HMM Profile Comparison

To assess how well generated sequences capture the statistical properties of a target protein family, we compare Hidden Markov Model (HMM) profiles built from generated sequences against profiles built from reference family alignments. The pipeline consists of three steps.

First, the generated sequences must be placed into the coordinate frame of the reference family MSA before any column-wise comparison is meaningful. We use MAFFT [23] to insert generated sequences into the reference family MSA, while preserving the original alignment length and prevent the reference columns from being perturbed. This ensures that position *i* in the resulting alignment corresponds to the same residue position across both reference and generated sequences.

Second, we build two HMM profiles using the HH-suite [48]. The reference profile is constructed directly from the reference MSA. The generated profile is constructed from the aligned generated sequences extracted in Step 1. Each profile encodes per-column amino acid emission probabilities as a position-specific frequency matrix (PSFM), capturing the residue composition and evolutionary variation at each position in the alignment. A minimum coverage threshold is applied: columns where fewer than 30% of reference sequences contribute a non-gap residue are excluded, as such columns provide unreliable frequency estimates.

Third, the two profiles are compared using again the HH-suite [48]. This produces several complementary similarity metrics: the alignment probability (Prob), E-value, and bit score. Collectively, these reflect how structurally similar the two profiles are (i.e., the degree to which the generated sequences reproduce the column-wise amino acid distributions of the reference family). A high bit score, combined with a low E-value, indicates that the generated sequences faithfully capture the positional residue preferences of the target family.

#### A.8.4 Positional KL-divergence comparison

To quantify how closely the residue distributions of generated sequences match those of a reference protein family at each alignment position, we compute the column-wise Kullback–Leibler (KL) divergence between position-specific frequency matrices (PSFMs) derived from generated and reference sequences respectively. The pipeline consists of three steps.

First, as in the HMM profile comparison, generated sequences are first inserted into the reference MSA coordinate frame using MAFFT [23], ensuring that column indices are shared between the reference and generated alignments.

Second,a PSFM of shape (*L×* 20) is computed independently for the reference MSA and the aligned generated sequences, where *L* is the alignment length and 20 corresponds to the standard amino acid alphabet (i.e., gaps are excluded from the KL computation). For each column *j*, residues are counted across all sequences, gap characters are ignored, and a small uniform pseudocount *ϵ* = 10^−6^ is added to every amino acid bin before normalising to a probability distribution. This prevents undefined logarithm terms at unobserved amino acids without materially distorting well-sampled columns. Columns where fewer than 30% of reference sequences contribute a non-gap residue are excluded from the analysis, as such positions carry unreliable frequency estimates.

Third, for each retained column *j*, we compute the KL divergence from the generated distribution *P*_*j*_ to the reference distribution *Q*_*j*_:

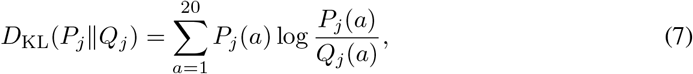

where the sum runs over the 20 standard amino acids. The mean KL divergence

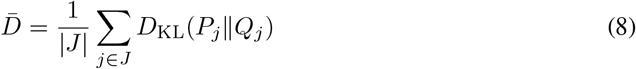

over all retained columns *J* provides a single scalar summary of how faithfully the generated sequences reproduce the positional residue preferences of the reference family. Lower values indicate closer agreement with the reference statistics, with 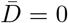 only when the generated and reference distributions are identical at every retained position.

### A.9 Sampling of held-out validation MSAs for conditional protein generation comparison

To ensure the validation set spans a broad range of MSA depths, we employed a stratified log-spaced sampling procedure. All eligible MSAs were first filtered to a depth range of 100 to 1000 sequences, and the resulting pool was partitioned into five logarithmically-spaced bins spanning that range (see Table 3). A fixed number of MSAs were then drawn uniformly at random from each bin, with any shortfall in underpopulated bins redistributed to the remaining ones. This procedure guarantees that the sampled validation MSAs are evenly distributed across depth magnitudes, from shallow MSAs with 100 sequences to deeper ones approaching 1000, avoiding the bias toward high-depth MSAs that would result from naive uniform sampling over the full pool.

**Table 3:**
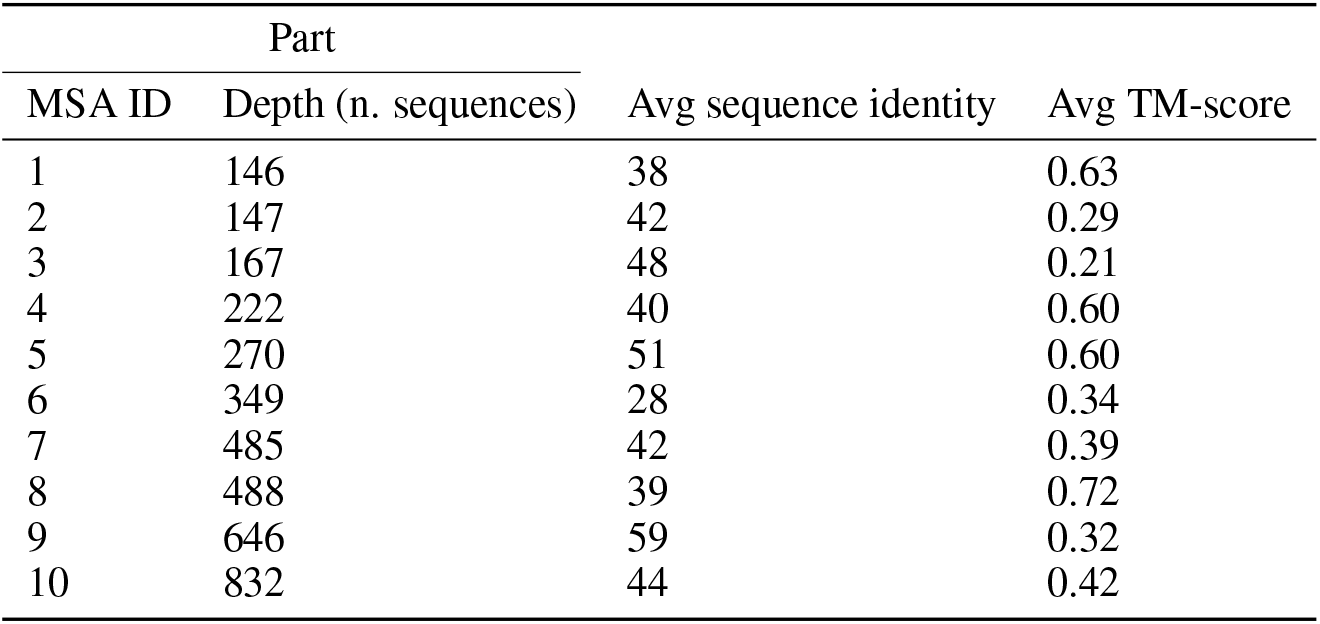
Statistics for 10 sampled held-out validation MSAs.

| Part |  | Avg sequence identity | Avg TM-score |
| --- | --- | --- | --- |
| MSA ID | Depth (n. sequences) |  |  |
| 1 | 146 | 38 | 0.63 |
| 2 | 147 | 42 | 0.29 |
| 3 | 167 | 48 | 0.21 |
| 4 | 222 | 40 | 0.60 |
| 5 | 270 | 51 | 0.60 |
| 6 | 349 | 28 | 0.34 |
| 7 | 485 | 42 | 0.39 |
| 8 | 488 | 39 | 0.72 |
| 9 | 646 | 59 | 0.32 |
| 10 | 832 | 44 | 0.42 |

### A.10 Conditional generation comparison for PDB MSAs

Here, we evaluate SFT versus few-shot prompting of ProtGPT3-MSA across 10 MSAs randomly sampled from the Protein Data Bank subset of OpenProteinSet [1] (see Table 4). Because each family in this subset is anchored to experimentally determined structures, it provides a more structurally consistent evaluation setting than the held-out validation MSAs tested above, where reference quality can vary across families. This makes it a complementary benchmark for assessing whether the trends observed on the validation set hold on families with higher-confidence structural annotations, albeit without guarantees of non-overlap with the MSA model training data. Each MSA was treated as an independent evaluation instance. Similarity to previous evaluations, we constructed a train/validation split for each sampled MSA, and used the training split both to fine-tune ProtGPT3-112M and to prompt ProtGPT3-MSA. We then generated 90 sequences per model (10 sequences for each combination of top-p ∈ {0.7, 0.9, 1.0} and temperature ∈ {0.8, 0.9, 1.0}). Finally, for each MSA, we compared the generated sequences from both models against the corresponding held-out validation split (i.e., sequences not used for either SFT or prompting) using metrics of sequence-, structure-, and family-level consistency (Appendix A.8). Fig 16 shows prompting ProtGPT3-MSA outperforms SFT of ProtGPT3-112M by generating sequences that more closely reflect the target protein family across all measured metrics, despite ProtGPT3-MSA not having been trained on those families. To get a concrete measure of how well ProtGPT3-MSA does on the family-level consistency metrics (i.e., HHalign and positional KL-divergence), we included two baselines, “Bootstrap MSA” and “Scrambled MSA”, respectively approximating upper and lower bound performance on these metrics. Specifically, Bootstrap MSA sample 30% of each true MSA sequences, performing the analysis for sequences that come from the true family distribution for each MSA. Conversely, “Scrambled MSA” provides a negative control, by running the family-level consistency analysis for ProtGPT3-MSA generated sequences for each MSA against a different MSA. This ensures the generated sequences by ProtGPT3-MSA do not just match per-column statistics for any MSA. Fig 16d-f shows ProtGPT3-MSA generated sequences achieve family-level consistency metrics close to sequence truly sampled from those family across most tested MSAs (i.e., Bootstrap MSA).

**Table 4:** Statistics for 10 sampled validation PDB MSAs Part.

| Part |  |  |  |
| --- | --- | --- | --- |
| MSA ID | Depth (n. sequences) | Avg sequence identity | Avg TM-score |
| 1 | 294 | 33.0763 | 0.795483 |
| 2 | 297 | 38.1639 | 0.868478 |
| 3 | 294 | 38.8053 | 0.719352 |
| 4 | 246 | 32.8433 | 0.826385 |
| 5 | 298 | 61.7558 | 0.916156 |
| 6 | 295 | 33.7895 | 0.840134 |
| 7 | 242 | 40.4272 | 0.797938 |
| 8 | 201 | 56.7782 | 0.715537 |
| 9 | 81 | 41.0078 | 0.762180 |
| 10 | 298 | 33.2846 | 0.654797 |

**Figure 15:**
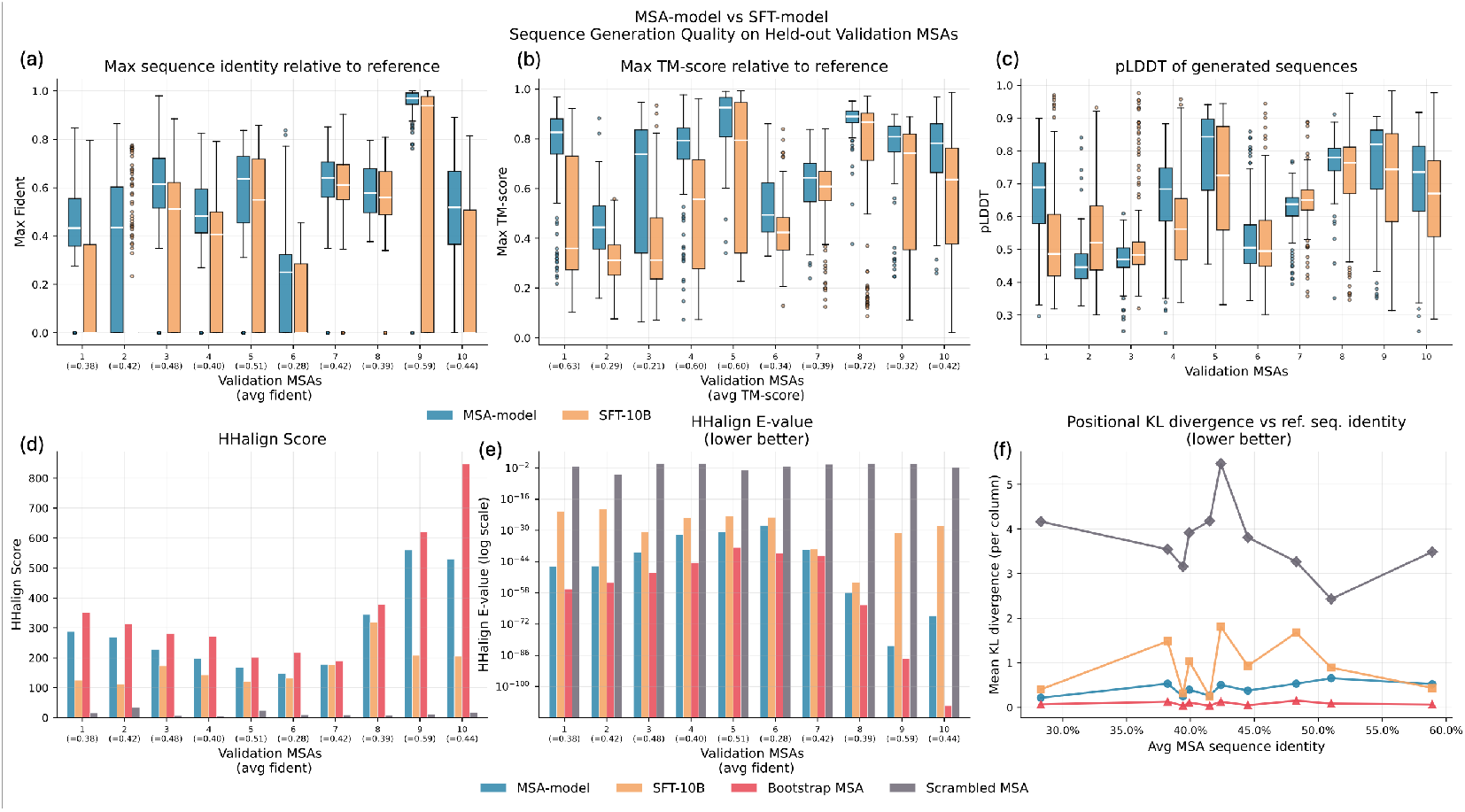
Conditional generation comparison prompting ProtGPT3-MSA vs SFT (single-sequence) ProtGPT3-10B for 10 held-out validation MSAs from ProtGPT3-MSA pre-training validation set.

**Figure 16:**
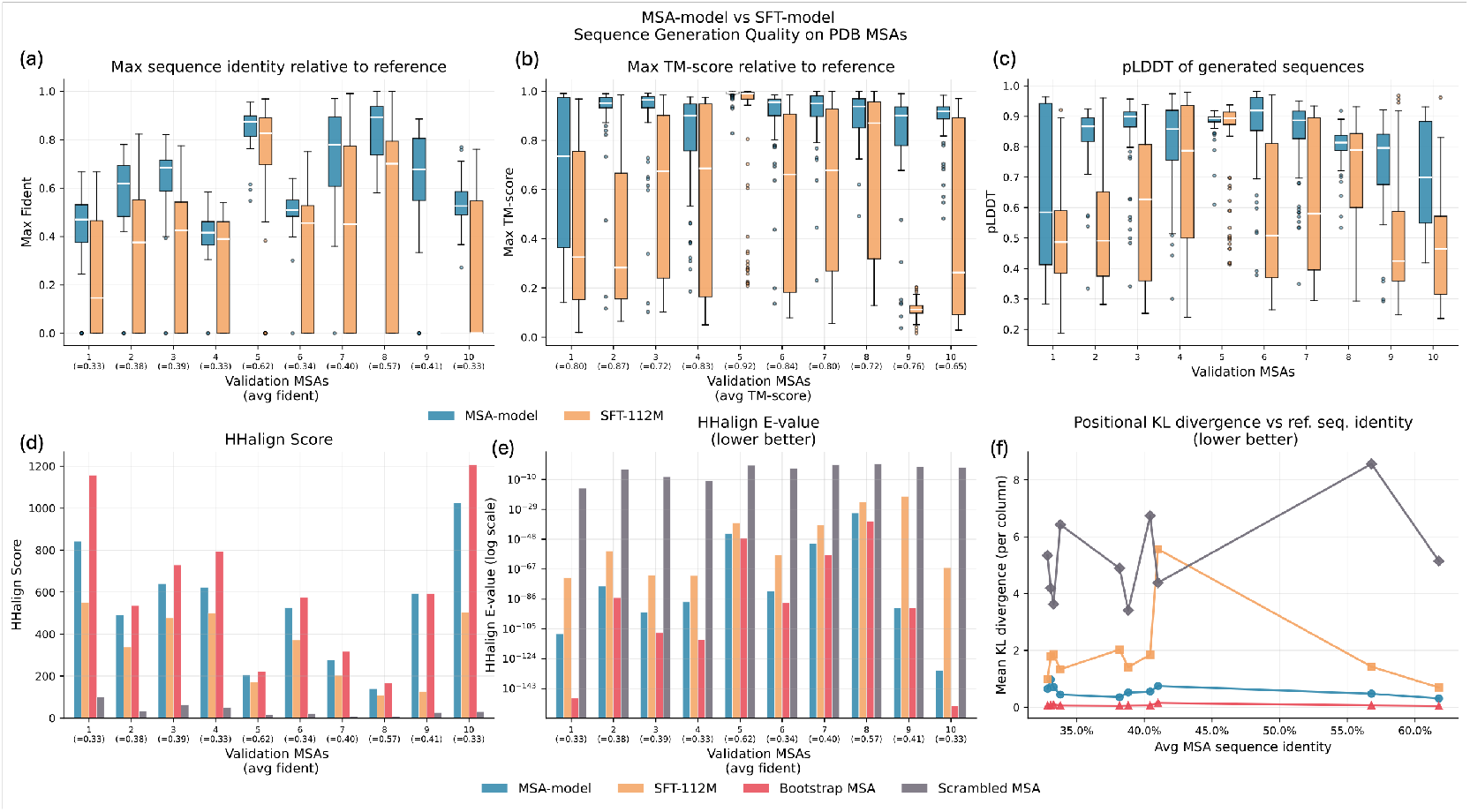
Conditional generation comparison prompting ProtGPT3-MSA vs SFT (single-sequence) ProtGPT3-112M for 10 randomly sampled MSAs from the Protein Data Bank subset of OpenProtein-Set [1]

**Figure 17:**
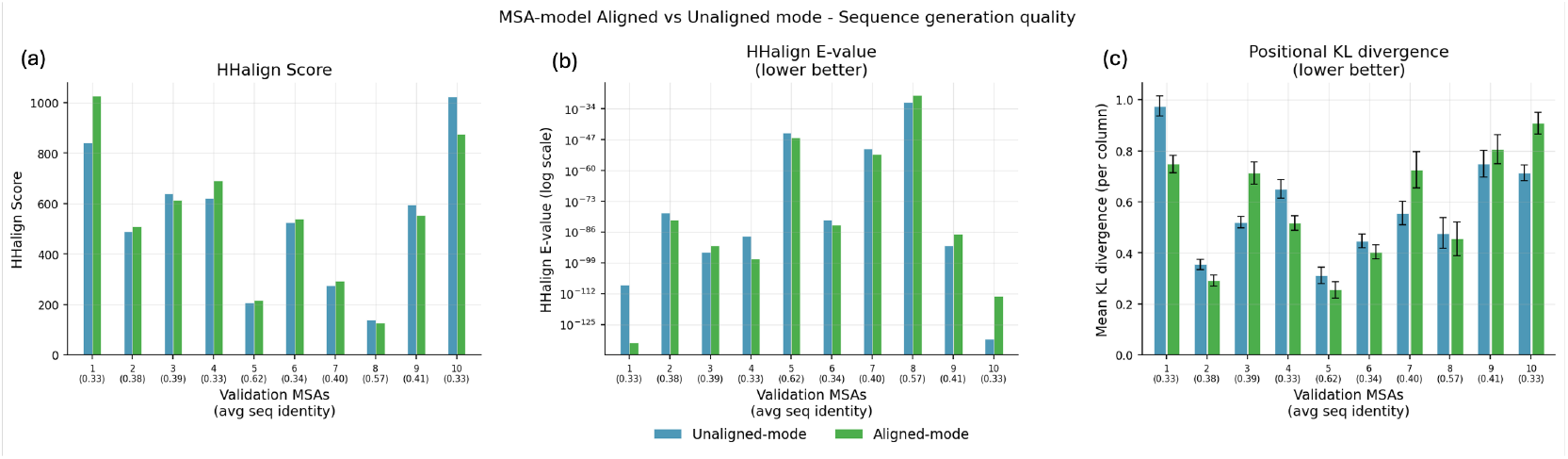
Comparison between sequences generated by ProtGPT3-MSA in *aligned* vs *unaligned* mode in terms of (a-b) HHM profile and (c) per-column KL-divergence relative to 10 reference MSAs. Within brackets on the x-axis, we report the average sequence identity for each reference MSA.

**Figure 18:**
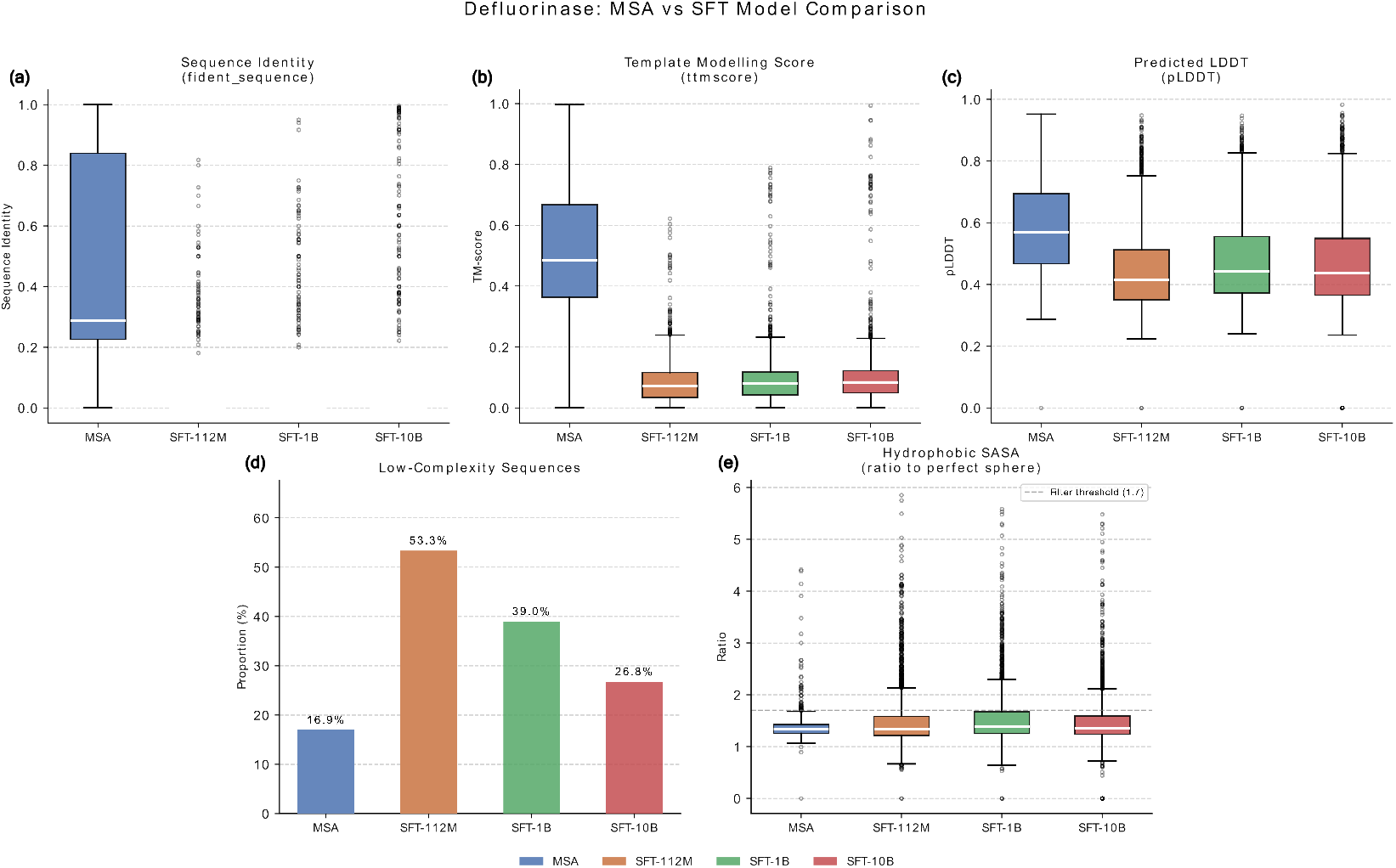
Computational characterization of defluorinases generated with ProtGPT3-MSA and single-sequence models prompted/fine-tuned with the available 7 sequences of experimentally confirmed defluorinases.

**Figure 19:**
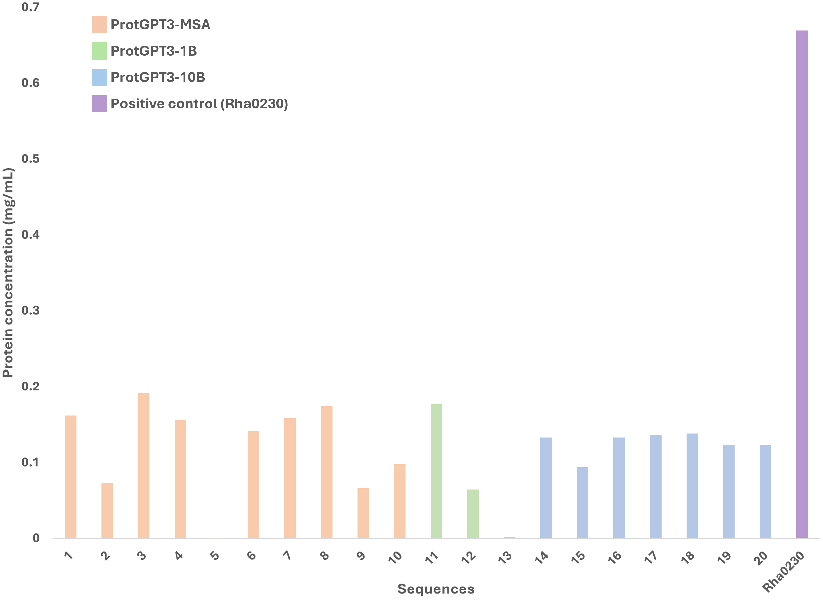
Protein concentration of the designed enzymes and the positive control Rha0230 after expression and purification. Sequences 1 to 10 were designed with ProtGPT3-MSA (salmon), 11 to 13 with ProtGPT3-1.3B (light green), and 14 to 20 with ProtGPT3-10B (mauve). Positive control Rha0230 from *R. jostii* is depicted in purple. Please note that sequences #5 and #13 show no protein concentration value since their cloning was not achieved

**Figure 20:**
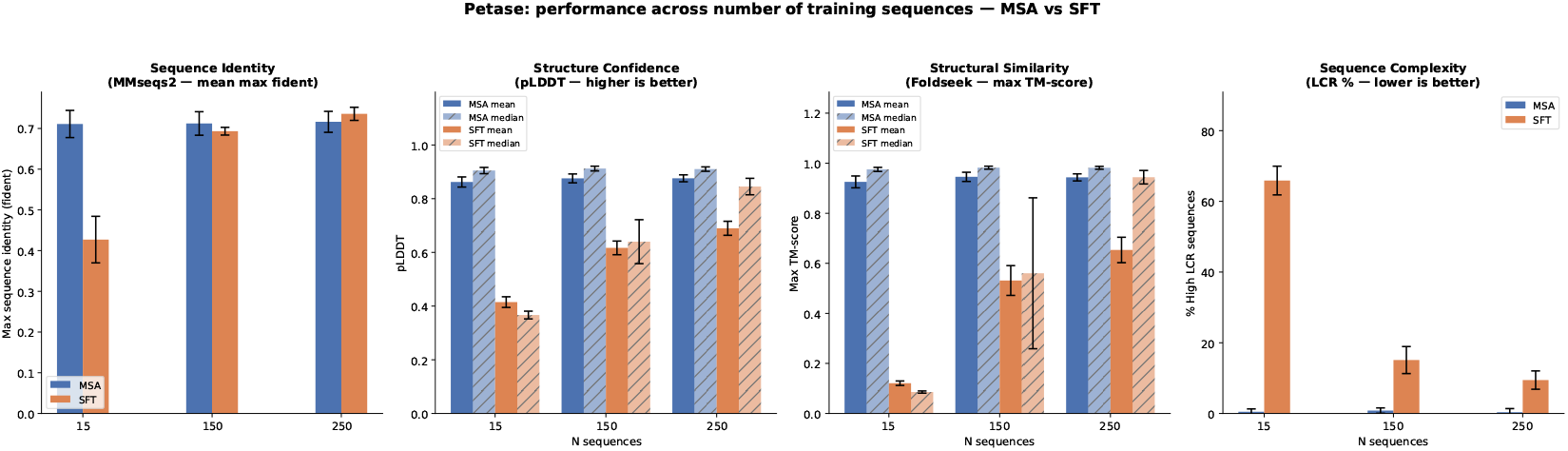
Comparison between sequences generated by few-shot prompting ProtGPT3-MSA vs SFT ProtGPT3-112M on Cutinase enzymes across different numbers of homologs (15, 150, 250).

**Figure 21:**
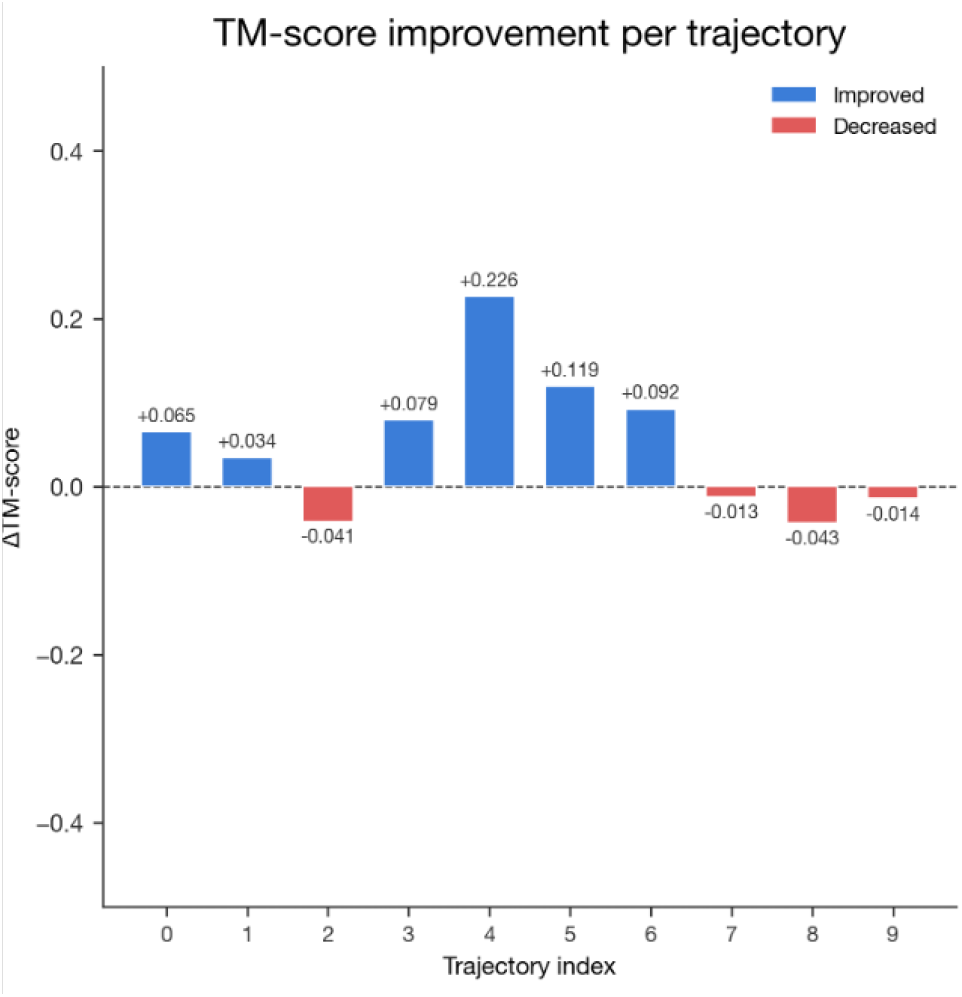
Change in TM-score per tested pairs of homologs after applying Feynman–Kac-style sequential Monte Carlo in ESM2 embedding distance. While ESM2 cosine similarity and TM-score are not expected to correlate directly [35], several pairs of homologs nonetheless show a modest increase in TM-score, suggesting that embedding-space steering can carry some structural signal.

**Figure 22:**
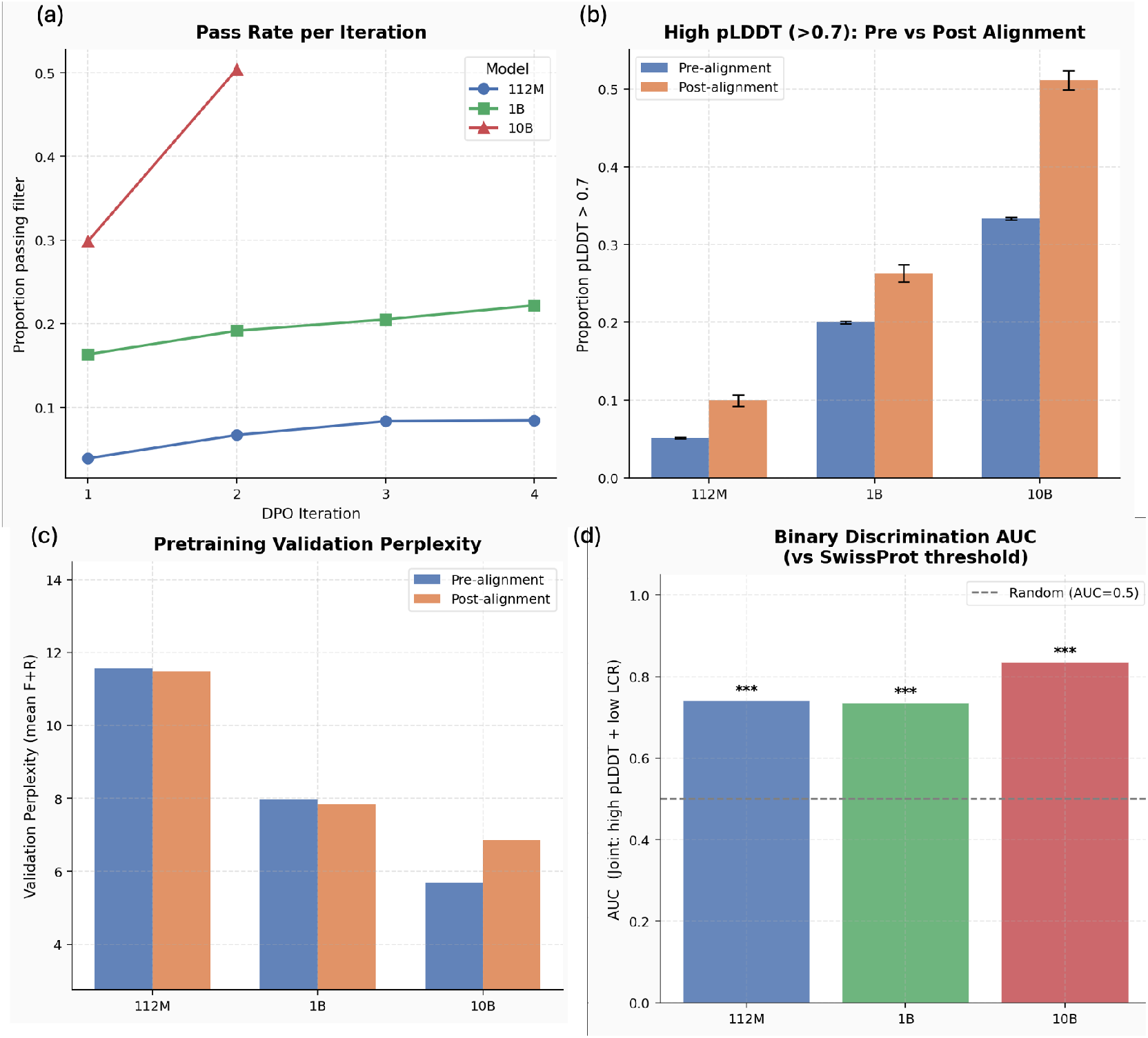
a) Pass rate denotes the proportion of generated sequences with plDDT *>* 0.7 as well as percentage of low complexity regions *<* 25%, which grows at each iteration of the DPO alignment pipeline. b) Proportion of sequence with plDDT *>* 0.7 before and after alignment. c) Model perplexity on the held-out pretraining validation sequences before and after alignment, which can be used as a proxy for model forgetting pretraining during the DPO alignment pipeline. d) AUC-ROC for each model intrinsic reward based on true sequences randomly sampled from the PDB dataset.

### A.11 Aligned vs Unaligned conditional protein generation with ProtGPT3-MSA

ProtGPT3-MSA was trained in to main modalities 1) *aligned*: the 16 input sequences were passed to the model aligned including gap tokens (i.e., “ − “) and 2) *unaligned*: the 16 input sequences were passed in full (i.e., including insertions and without gaps). Therefore, the model has the ability of generating both *aligned* and *unaligned* protein sequences. When we compare few-shot prompting ProtGPT3-MSA with SFT of single-sequence ProtGPT3 models, we always used the *unaligned* modality to ensure a fair comparison (i.e., single-sequence models were not trained to process aligned sequences). Here, we investigate any potential difference in homolog generation quality between the two modalities. In particular, we look at potential difference in the quality of generated sequences at the level of MSA per-column statistics to assess any potential benefit of inputting already aligned sequences to the model. For this comparison, we randomly sampled 10 MSAs from the Protein Data Bank subset of OpenProteinSet [1]. From each MSA, we randomly sampled a set of 15 homologs and used it to (few-shot) prompt ProtGPT3-MSA in both *aligned* and *unaligned* mode. Next, we compared the quality of the generated sequence from each modality relative to the reference MSA using the positional KL-divergence and HMM profile comparison described in Appendix A.8. Fig. 17 shows there is not substantial difference in the quality of generated sequences between the two modalities.

### A.12 Conditional generation comparison for defluorinases

#### Design and Computational Characterization

ProtGPT3-MSA and single-sequence models were used for the targeted generation of defluorinases. Sequences with ProtGPT3-MSA were generated by prompting with the 7 homologous sequences reported in Probst et al. [36]. We used the same 7 sequences to fine-tune the three single-sequence ProtGPT3 models. We used k-fold cross validation to pick the optimal number of epoch to train each model. Next, we generated 3240 sequences per model (360 sequences for each combination of top-p ∈ {0.7, 0.9, 1.0} and temperature ∈ {0.8, 0.9, 1.0}) and predicted their homodimeric structures with Boltz2 [34] in single-sequence mode, while providing a template pdb file of an experimentally solved crystal structure from a defluorinase (PDB: 3UMG_A). All subsequent structural analyzes were performed with the highest-ranking structure of the 5 generated predictions. Structural and sequential similarity comparisons against the reference protein structure and confirmed 7 defluorinase sequences were performed as explained above (see A.8). Additionally, we used PyRosetta [7] to compute Rosetta Energy Scores [26, 3], and the ratio between the hydrophobic solvent accessible area of the protein complex and the idealized surface area for a protein complex of the same size as described by [12]. For computational characterization, we define a broad success metric that classifies a design as successful if the generated sequence resembles a defluorinase sequentially (sequence identity to any known defluorinases between 85% and 95%), structurally (TM-Score ≥ 0.5) and passes a mild solubility filter (hydrophobic solvent accessible solvent area ratio ≤ 1.7). Given the unknown molecular features that separate defluorinases from dechlorinases, we reasoned that staying close in sequence space and structural space to experimentally confirmed defluorinases will be necessary to achieve reasonable experimental success rates in the absence of better filter criteria. Filtering for the hSASA ratio, is a high-level filter for protein solubility [55]. We note that these filters can only be considered a minimum requirement for a successful design and do not reflect experimental success rates, which are expected to be lower. However, as only very few sequences generated by the fine-tuned ProtGPT3-1.3B and ProtGPT3-10B passed this initial set of filters, further filtering was only possible with sequences generated with ProtGPT3-MSA. Specifically, we ranked sequences by their Boltz2 pLDDT value [34], spatial aggregation propensity, hSASA ratio, Rosetta energy score [7, 26, 3], interface pLDDT value, and the ligand-protein interface pTM score, hypothesizing that these would translate to higher experimental success rates beyond the initial broad filtering.

#### Cloning, expression and purification of dehalogenases

The encoding sequences of the 20 candidates and the *R. jostii* Rha0230 dehalogenase were purchased from GenScript Biotech^®^ as double stranded DNA with flanking recognition sites for *Bsa*I at both 5’and 3’ ends. Such sequences were cloned by Goldengate in the pCoofy plasmid. *E. coli* DH5*α* cells were then transformed by heat-shock with the Goldengate products and streaked onto LB-agar plates supplemented with kanamycin (50 *µ*g/mL) and 1 mM isopropyl-*β*-D-1-thiogalactopyranoside (IPTG), presence of the genes of interest was confirmed by colony PCR and the expression vectors isolated using the GeneJET Plasmid Miniprep Kit from ThermoFisher Scientific^®^ (Cat. K0502).

Protein expression of the designs and the positive control was carried out in *E. coli* BL21(DE3) cells, after transformation with the expression vectors by heat-shock, in 2YT auto-induction medium (auto2YT) purchased from Formedium™ (Cat. AIM2YT0260). Over-night (O/N) precultures were grown at 37ºC in 3 mL LB+kanamycin (50 *µ*g/mL), then 40 *µ*L were taken to inoculate 4 mL auto2YT+kanamycin/construct. These cultures were incubated at 37ºC for 4 hours in 24-well plates with mild agitation, then transferred to 20ºC O/N.

Cells were harvested by centrifugation (3000 g, 20 minutes, 4ºC) and resuspended in 1 mL of BUFFER A (50 mM TRIS·H_2_SO_4_, 500 mM NaAcO, 20 mM imidazole), supplemented with 1X cOmplete™ protease inhibitor cocktail EDTA-free from Merck (Cat. 11873580001) and a few crystals of DNAse and lysozyme. Cell lysis was triggered by the addition of 1X FastBreak™ Cell Lysis Reagent from Promega^®^Corporation (Cat. V8571) and incubation at room temperature (r.t.) for 30 minutes with mild mixing, then lysates were centrifuged (3000 g, 1 h, 4ºC) and the supernatants recovered for batch purification with Pierce™ Ni-NTA Magnetic Agarose Beads (ThermoFisher Scientific^®^, Cat. 78606). The purification protocol briefly consisted in one hour of incubation with the beads, removal of the supernatant, two wash steps with BUFFER B (50 mM TRIS·H_2_SO_4_, 500 mM NaAcO, 40 mM imidazole), and one single elution step with BUFFER C (50 mM TRIS·H_2_SO_4_, 500 mM NaAcO, 400 mM imidazole). Presence of the purified candidates and the positive control was checked by Bradford assay (Fig. 19), in which absorbance was recorded at 595 nm on a TECAN plate reader. Sequences #5 and #13 show no protein concentration value since these are the candidates whose cloning was not achieved. Such purification procedure was achieved in an Opentrons Flex liquid-handling robot.

### A.13 Conditional generation comparison for Cutinase enzymes

Cutinases were selected as a test case for targeted sequence generation. Cutinases (EC 3.1.1.74) are serine esterases in the *α*/*β*-hydrolase family that hydrolyze cutin, a waxy polyester that coats plant surfaces. They are produced by many phytopathogenic fungi and bacteria. Beyond their natural role, cutinases have attracted growing industrial interest because some members of this family can also hydrolyze polyethylene terephthalate (PET), a common plastic used in food packaging and polyester fibers, releasing terephthalic acid (TPA) and ethylene glycol. To evaluate targeted generation in this family, we prompted ProtGPT3-MSA with increasing numbers of cutinase homologs (15, 150, and 250 sequences) and compared its outputs with those of a single-sequence model, ProtGPT3-112M, that was supervised fine-tuned on the same subsets of sequences. Across all data regimes, the MSA-based model generated more plausible cutinase-like sequences than the fine-tuned single-sequence model, achieving higher sequence identity to the held-out validation set, higher structural confidence (pLDDT), and higher structural similarity (TM-score) (see Fig. 20). Although the single-sequence model improved as more training sequences were provided, reaching values close to the MSA model at 250 sequences for pLDDT and TM-score, the MSA model performed consistently well even with as few as 15 input homologs. Sequence and structure similarity as well as pLDDT were estimated as previously detailed (see Appendix A.8). The MSA model also produced sequences with substantially lower low-complexity content than the fine-tuned single-sequence model. Low-complexity regions (LCRs) are sequence segments with low amino-acid diversity or repetitive composition, identified using standard sequence-complexity filters (see Fig. 20 right-hand side panel). These results indicate the ability of the MSA model to generate realistic family-specific sequences even with limited input information.

### A.14 FK-SMC Sampling with ProtGPT3-MSA

#### Algorithm 1

FK-SMC Sampling with ProtGPT3-MSA

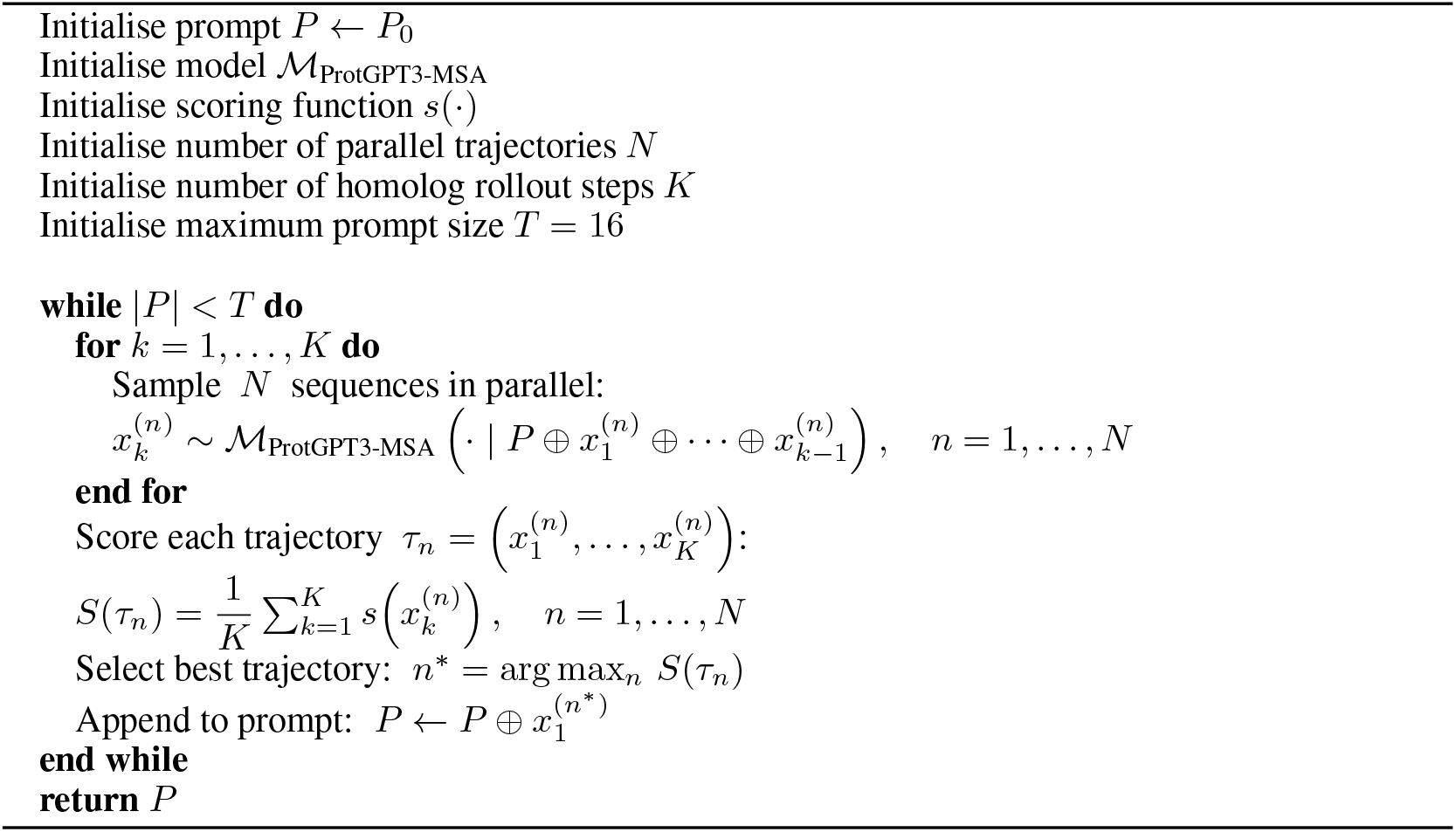

We adapt a Feynman–Kac-style sequential Monte Carlo (FK-SMC) procedure to steer sequence generation from ProtGPT3-MSA toward a target objective, *s*(·). The procedure operates iteratively: at each step, the model is prompted with a growing sequence of homologs and branches into multiple candidate trajectories, which are scored and selectively retained to bias the next generation step. Formally, let *P*_0_ denote an initial prompt of one or more concatenated homolog sequences, *N* the number of candidate trajectories sampled per step, and *K* the rollout length (number of homolog sequences per trajectory). At each step *t*, we sample *N* trajectories from ProtGPT3-MSA conditioned on the current prompt *P*_*t* 1_. Each trajectory 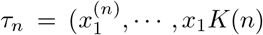 consists of *K* homolog sequences generated autoregressively. Trajectories are scored by averaging a sequence-level objective *s*(·) across all *K* sequences. The highest-scoring trajectory is selected, its first sequence is appended to the prompt, and the process repeats. Both *N* and *K* are controllable hyperparameters governing the breadth and depth of the search. In practice, we terminate the generation when the prompt reaches 16 sequences (i.e., the max n. of sequences that ProtGPT3-MSA can process), although the search could continue indefinitely by simply starting to remove the first sequence in the prompt at each new step. The procedure is summarised in Algorithm 1.

For the steering experiment in ESM2 embedding space (described in Section 5), we sample pairs of homologous sequences with TM-score *∈* [0.5, 0.6] from OpenProteinSet/PDB subsets [1], randomly designating one as the starting sequence and the other as the target. We use ESM2-150M [27] as the embedding model, extracting representations from the final layer, and computing cosine similarity to the target sequence as the scoring function *s*(·). We set the number of homolog rollout steps as *K* = 5 and the number of parallel trajectories *N* = 900, repeating the experiment across 10 randomly sampled homolog pairs. The *N* trajectories are sampled in parallel at each homolog rollout step, which is computationally feasible due to the relatively small size of ProtGPT3-MSA (112M parameters), allowing a large number of trajectories to be batched efficiently on a single GPU. We implement custom inference code to support this parallel sampling scheme, with the additional option of caching the key-value (KV) representations of the prompt at each planning iteration, amortising the cost of re-encoding the growing prompt across trajectories at the expense of additional memory cost. We release this code alongside the models; the implementation is compatible with any scoring function defined over individual protein sequences, making it straightforward to adapt to alternative objectives beyond cosine similarity in embedding space.

### A.15 Single-sequence model alignment to high complexity and high pLDDT

To improve the quality of generated sequences across the 3 sizes of single-sequence models, we generate 80 thousands sequences across all combinations of top-p 0.6, 0.8, 0.9, 1.0 and temperature 0.6, 0.8, 0.9, 1.0 for each model. Next, we score all generated sequences based on % of low complexity regions as well as pLDDT (predicted by ESMFold [27]) and classify them as “pass”, each sequence that has plDDT *>* 0.7 as well as percentage of low complexity regions *<* 25%, while classifying all other sequences as “fail”. Next, we cluster “pass” and “fail” sequence separately at 50% sequence identity and 0.8 coverage using MMseqs2 [47]. We take a single sequence form each cluster and use them to build a binary preference dataset. Since the ratio of “fail” sequence was much higher than the “pass” one, especially for the smaller models, we build the preference dataset by sampling 20 “fail” sequences for each “pass” sequence. Importantly, we enforce a strict sequence length match between pairs of “pass” and “fail” sequences to avoid the model learning to distinguish the two in terms of sequence length instead of complexity and pLDDT. Based on this preference dataset, we use the Direct Preference Optimization (DPO) [38] algorithm to shift each model distribution towards high complexity and high pLDDT sequences. We repeat this entire process for 3 iteration, apart from ProtGPT3-10B, which after a single iteration achieved a 5% proportion of low complexity sequences.

#### A.15.1 Intrinsic Reward Discrimination Analysis

DPO alignment was performed using model-generated sequences scored against pLDDT and LCR thresholds; here we test whether the aligned models have internalised these objectives when evaluated on real, naturally occurring protein sequences. We randomly subsample sequences from the PDB, which include AlphaFold-predicted pLDDT scores, and compute the percentage of low-complexity regions (LCR) for each sequence. Sequences are partitioned into two groups using the thresholds applied during DPO training: *low-quality* sequences have pLDDT *<* 70 and percentage of low complexity regions *>* 25%, and *high-quality* sequences have pLDDT ≥ 70 and percentage of low complexity regions *<* 25%, with groups balanced by random subsampling of the majority class. For each aligned model (112M, 1.3B, 10B parameters), we compute the intrinsic reward of each sequence relative to its corresponding base model,

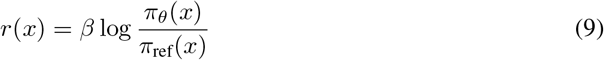

where *π*_*θ*_ is the aligned model, *π*_ref_ is the base (reference) model and *β* is the KL penalty coefficient use in DPO. We report the AUC-ROC of *r*(*x*) as a binary classifier between the two groups, with a t-test for significance (see Fig. 22d). AUC values in the range 0.7–0.8 across model sizes indicate that the aligned models assign systematically higher intrinsic rewards to high-quality sequences, demonstrating generalisation of the learned preference to real protein sequences not seen during alignment. Importantly, we do not expect nor desire a near-perfect AUC: low-pLDDT and high-complexity sequences are naturally occurring and present in the pretraining corpus, so both *π*_*θ*_ and *π*_ref_ assign non-trivial likelihood to them. Since *r*(*x*) is a relative quantity, a randomly drawn low-quality sequence may still receive a higher intrinsic reward than a randomly drawn high-quality sequence if it is intrinsically more likely under the base model (e.g., short or repetitive sequences that are statistically common). A perfect AUC would indicate the model has learned to categorically reject an entire class of naturally occurring proteins, which would reflect overfitting to the alignment objective rather than a balanced internalisation of structural preferences.

### A.16 Hyperparameters

We report the main hyperparameters across the 4 models in the tables below, all other hyper-parameters can be found in the model configuration and code at https://huggingface.co/collections/AI4PD/protgpt3-family and https://github.com/michele1993/protGPT3.

**Table 5:** Hyperparameters used to train the ProtGPT3-MSA.

| Hyperparameter | Value |
| --- | --- |
| Max sequence length | 16,384 |
| Router aux. loss coef. | 0.05 |
| Optimizer | AdamW |
| Learning rate | $2 \times 10^{-4}$ |
| $(\beta_1, \beta_2)$ | (0.9, 0.95) |
| Weight decay | 0.1 |
| Gradient clipping | 1.0 |
| Grad. accum. steps | 16 |
| Max tokens per batch | 100,000 |
| Precision | bfloat16 |
| N. training GPUs | 4 |

**Table 6:** Hyperparameters used to train the ProtGPT3-112M.

| Hyperparameter | Value |
| --- | --- |
| Max sequence length | 1024 |
| Router aux. loss coef. | 0.05 |
| Optimizer | AdamW |
| Learning rate | $5 \times 10^{-4}$ |
| $(\beta_1, \beta_2)$ | (0.9, 0.999) |
| Weight decay | 0.1 |
| Gradient clipping | 1.0 |
| Grad. accum. steps | 1 |
| Batch size | 500 |
| Precision | bfloat16 |
| N. training GPUs | 4 |

**Table 7:** Hyperparameters used to train the ProtGPT3-1.3B.

| Hyperparameter | Value |
| --- | --- |
| Max sequence length | 1024 |
| Router aux. loss coef. | 0.05 |
| Optimizer | AdamW |
| Learning rate | $2.5 \times 10^{-4}$ |
| $(\beta_1, \beta_2)$ | (0.9, 0.999) |
| Weight decay | 0.1 |
| Gradient clipping | 1.0 |
| Grad. accum. steps | 4 |
| Batch size | 100 |
| Precision | bfloat16 |
| N. training GPUs | 16 |

**Table 8:**
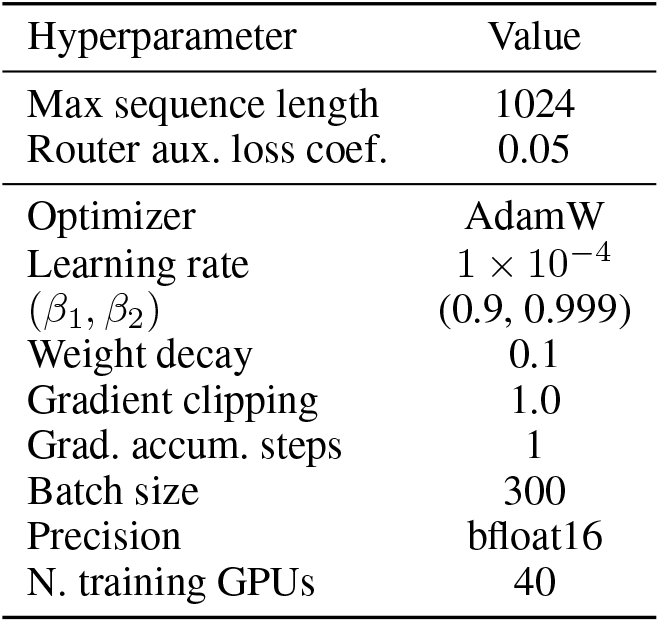
Hyperparameters used to train the ProtGPT3-10B.

### A.17 Infrastructure and optimization

All models were trained on NVIDIA H100 GPUs, using FlashAttention-2 [11], online mini-batch packing [25] as well as Liger Kernel [19] for maximal efficiency. Additionally, we used the Deep-Speed Library [41] for efficient distributed data parallel (DDP) across all models as well as for ZeRO-3 [39] for training, SFT and aligning the 10B model.

### A.18 Models and code

All 7 models are publicly available open source on Hugging Face Hub at https://huggingface.co/collections/AI4PD/protgpt3-family.

The entire code base for training and evaluation is available on a GitHub repository at https://github.com/michele1993/protGPT3.

